# Radiation survival selects clonally restricted NSCLC survivors with impaired metastatic capacity and distinct bioenergetic states

**DOI:** 10.64898/2026.11.18.767282

**Authors:** Marina Martinez-Cruz, Mohammad Darbalaei, Anna Baginska, Sven T Liffers, Viyan Arab, Vuk Đinović, Jennifer Jung, Johann Matschke, Verena Jendrossek, Alexander Schramm, Daniel Hoffmann, Barbara M. Grüner

## Abstract

Radiation resistance and recurrence remain major challenges in non–small cell lung cancer (NSCLC). Whether radiation survival enriches for more aggressive, metastasis-competent cells remains unresolved. Using longitudinal cellular barcoding, functional assays, and multi-omics profiling in diverse NSCLC models, we show that fractionated irradiation imposes a strong clonal bottleneck, but recovery follows either stochastic or deterministic trajectories. Despite this diversity, all radiation-surviving populations exhibit increased radiosensitivity and profoundly impaired metastatic capacity, including near-complete failure in lung colonization *in vivo*. Transcriptomic and metabolic analyses reveal three distinct bioenergetic states, hypermetabolic, energy-conserving, and metabolically suppressed, each supported by unique transcriptional programs. Crucially, clonal dominance does not predict molecular or functional state, indicating that clonal selection and post-irradiation reprogramming are decoupled. These findings challenge the assumption that radiation selects for aggressive clones and reveal that radiation survival may instead create a state of post-irradiation vulnerability. This has important implications for therapy: targeting adaptive pathways, not just resistant clones, may improve outcomes.

**STATEMENT OF SIGNIFICANCE:** This study challenges the long-standing assumption that radiation survival selects for more aggressive, metastasis-competent tumor cells. Instead, we show that radiation triggers a clonal bottleneck that leads to diverse molecular and metabolic trajectories, converging on a state of increased radiosensitivity and impaired metastatic fitness. Crucially, clonal selection and cellular adaptation are decoupled, revealing a previously unrecognized vulnerability. These findings suggest that radiation may inadvertently suppress metastatic spread, opening new therapeutic opportunities by targeting adaptive pathways in survivors.

Graphical Abstract:
Radiation survival selects clonally restricted NSCLC survivors with impaired metastatic capacity and distinct bioenergetic states.
Created in BioRender. Martínez-Cruz, M. (2027) https://BioRender.com/9u5o898

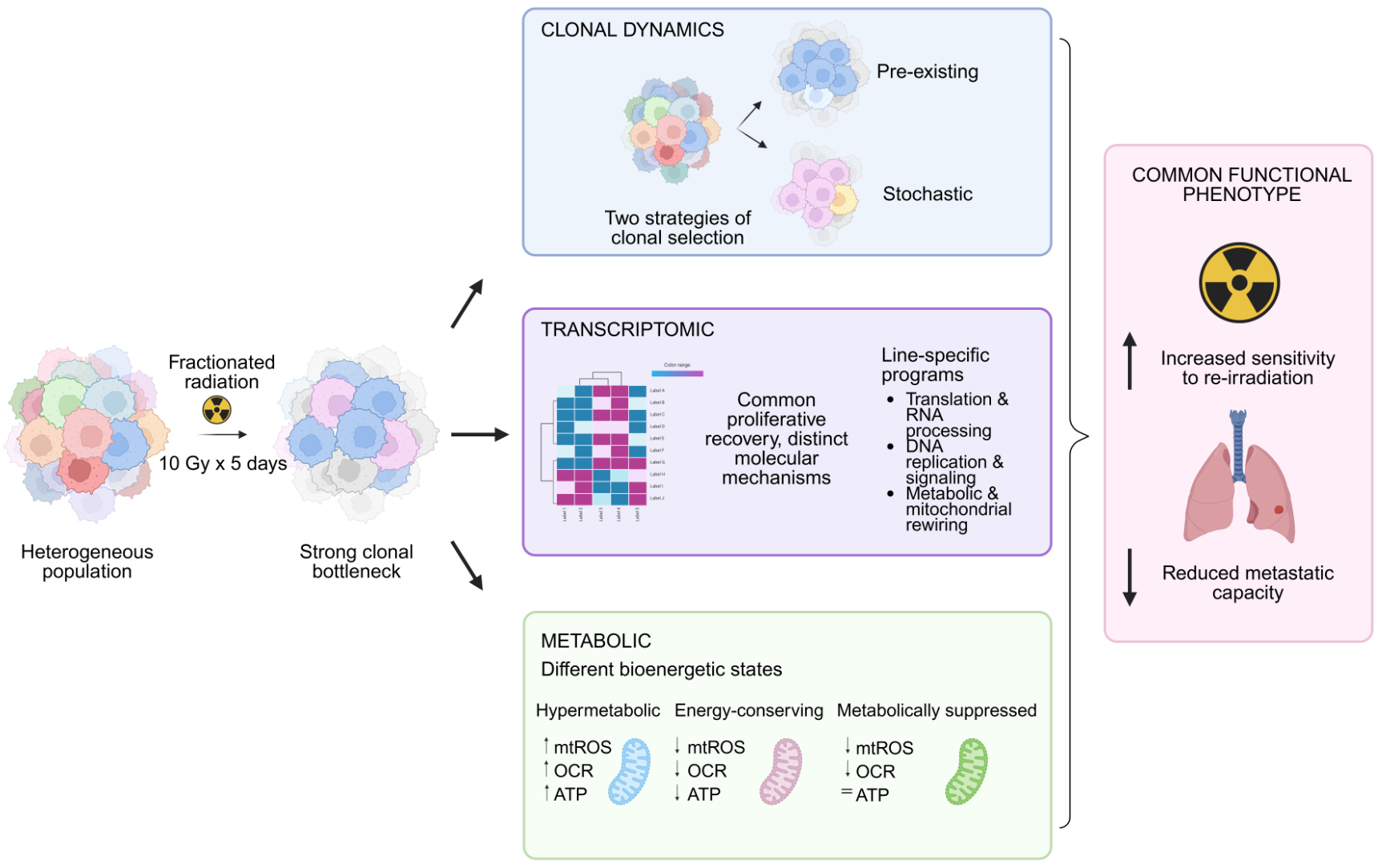

## INTRODUCTION

Lung cancer remains the leading cause of cancer-related mortality worldwide, with non-small-cell lung cancer (NSCLC) accounting for approximately 85% of all lung cancer cases (1). Despite advances in diagnosis and treatment, clinical outcomes remain poor, particularly in advanced disease, with metastatic dissemination being a major determinant of patient survival (2). NSCLC is characterized by extensive inter- and intratumor heterogeneity, which is reflected in differences in tumor evolution, treatment response, and metastatic behavior (2, 3). This heterogeneity complicates the identification of determinants of treatment response and raises the possibility that distinct tumor cell populations may differ not only in their response to radiation but also in their capacity for metastatic dissemination and subsequent adaptation (3).

Radiation therapy (RT) is an established component of NSCLC treatment and can be administered with curative purpose in localized disease and in selected patients with limited metastatic burden. However, its efficacy is limited by tumor radioresistance and by the toxicity associated with increasing radiation exposure (4). Ionizing radiation induces cellular stress primarily through DNA damage, particularly DNA double-strand breaks, while also promoting oxidative stress and altering mitochondrial and metabolic homeostasis (5, 6). Although these effects contribute to radiation-induced cell death, they may also influence the fate and adaptation of cells that survive treatment.

Importantly, tumor heterogeneity provides a potential basis for differential responses to irradiation. Tumors contain cellular subpopulations with distinct intrinsic radiosensitivities. Hence, repeated exposure to fractionated irradiation may impose a selective pressure favoring the survival and expansion of more resilient clones. Consistent with this model, previous studies have demonstrated substantial heterogeneity in radiosensitivity within individual tumors and have implicated clonal selection in the emergence of radiation-surviving populations (7, 8). However, the extent to which radiation survival reflects pre-existing clonal properties versus acquired adaptation, the mechanisms that sustain long-term survival, and the relationship between radiation survival and metastatic potential remain incompletely understood.

Here, we investigated the cellular and molecular basis of radiation survival in heterogeneous NSCLC populations and its relationship with metastatic potential. Using high-resolution DNA barcoding, transcriptomic profiling, and functional metabolic assays, we first examined whether radiation survival reflected pre-existing clonal differences or emerged as an acquired phenotype following radiation exposure. We then characterized the transcriptional and metabolic programs associated with long-term radiation survival and determined how radiation reshaped clonal composition over the course of treatment. Finally, we evaluated whether radiation survival was linked to changes in metastatic potential. Together, this approach allowed us to examine radiation survival as a dynamic process shaped by clonal selection and cellular adaptation.

## RESULTS

### Radiation survival imposes a clonal bottleneck with divergent recovery trajectories

To study the clonal evolutionary processes in heterogeneous NSCLC, we utilized six polyclonal cell lines established from either primary tumors or metastases of the well-established *Kras^G12D^;p53^KO^* genetically engineered mouse model (9). First, we characterized the radiation response of these NSCLC cells, using a clonogenic survival assay following radiation (10). All cell lines showed a dose-dependent decline in the surviving fraction, but the extent of this response varied markedly between lines. Notably, metastasis-derived cell lines tended to be more radiosensitive than primary tumor-derived lines, especially at higher doses (**Figure 1A**). These differences in radiosensitivity prompted us to investigate the clonal and molecular determinants of these responses, how they evolve under prolonged irradiation, and how they affect metastatic potential. To address this, we implemented a lentiviral cellular barcoding strategy (sCORE) (**Figure 1B**) that enables longitudinal tracking of single cells through stable barcode integration. To ensure the robustness of our system, we confirmed successful barcode labeling (>90% GFP^+^ after sorting (**Supplementary Figure S1A–D**)). Furthermore, we verified that barcode introduction had no detectable effect on cell proliferation and only minimal effects on radiosensitivity, with a dose-dependent difference observed in 482N1 (**Supplementary Figure S1B**). These results supported the use of the sCORE system for longitudinal tracking of barcode dynamics.

**Figure 1.**
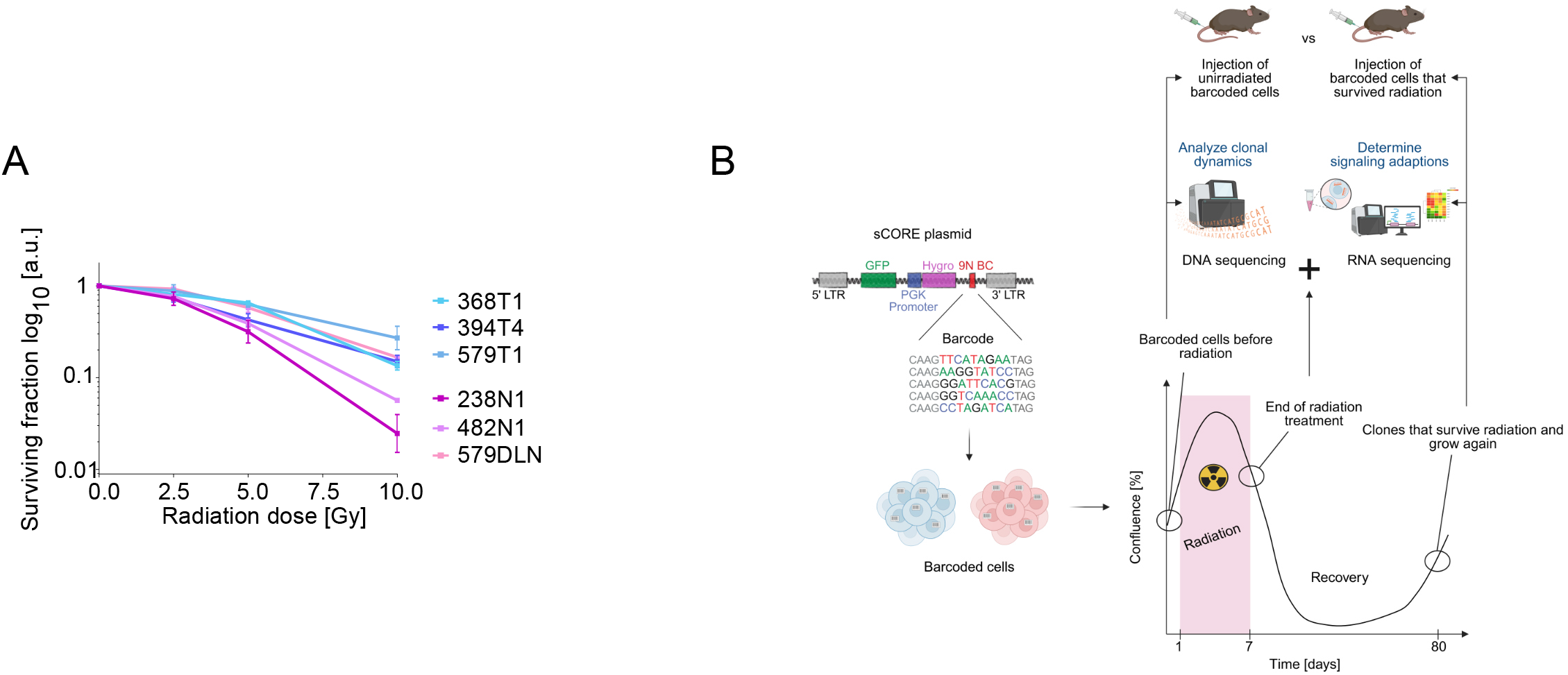
Heterogeneous radiosensitivity and clonal tracking setup. **(A)** Colony formation assay of the indicated murine NSCLC cell lines. Primary tumor-derived cell lines (368T1, 394T4, 579T1) are depicted in blue tones, whereas metastasis-derived cell lines (238N1, 482N1, 579DLN) are depicted in pink tones. Data are presented as Mean ± SEM from N=3 independent biological replicates. **(B)** Schematic of the experimental workflow. NSCLC cell lines were transduced with the sCORE barcoding system for clonal tracking. After expansion, cells were subjected to fractionated irradiation (5 × 10 Gy over 7 days). Genomic DNA and RNA were collected at multiple time points (T1–T4) for barcode and transcriptomic analysis. Surviving populations were expanded for *in vivo* metastatic assays. Created in BioRender. Martínez-Cruz, M. (2027) https://BioRender.com/vw2zbt3

Having confirmed the neutrality of the barcoding system, we next investigated the responses and clonal dynamics of NSCLC cells following fractionated irradiation (**Figure 2 and Supplementary Figure S2**). For this purpose, four barcoded cell lines (394T4, 482N1, 579T1, and 579DLN), together with two non-barcoded lines (368T1, 238N1), were subjected to this regimen (**Figure 2A**).

**Figure 2.**
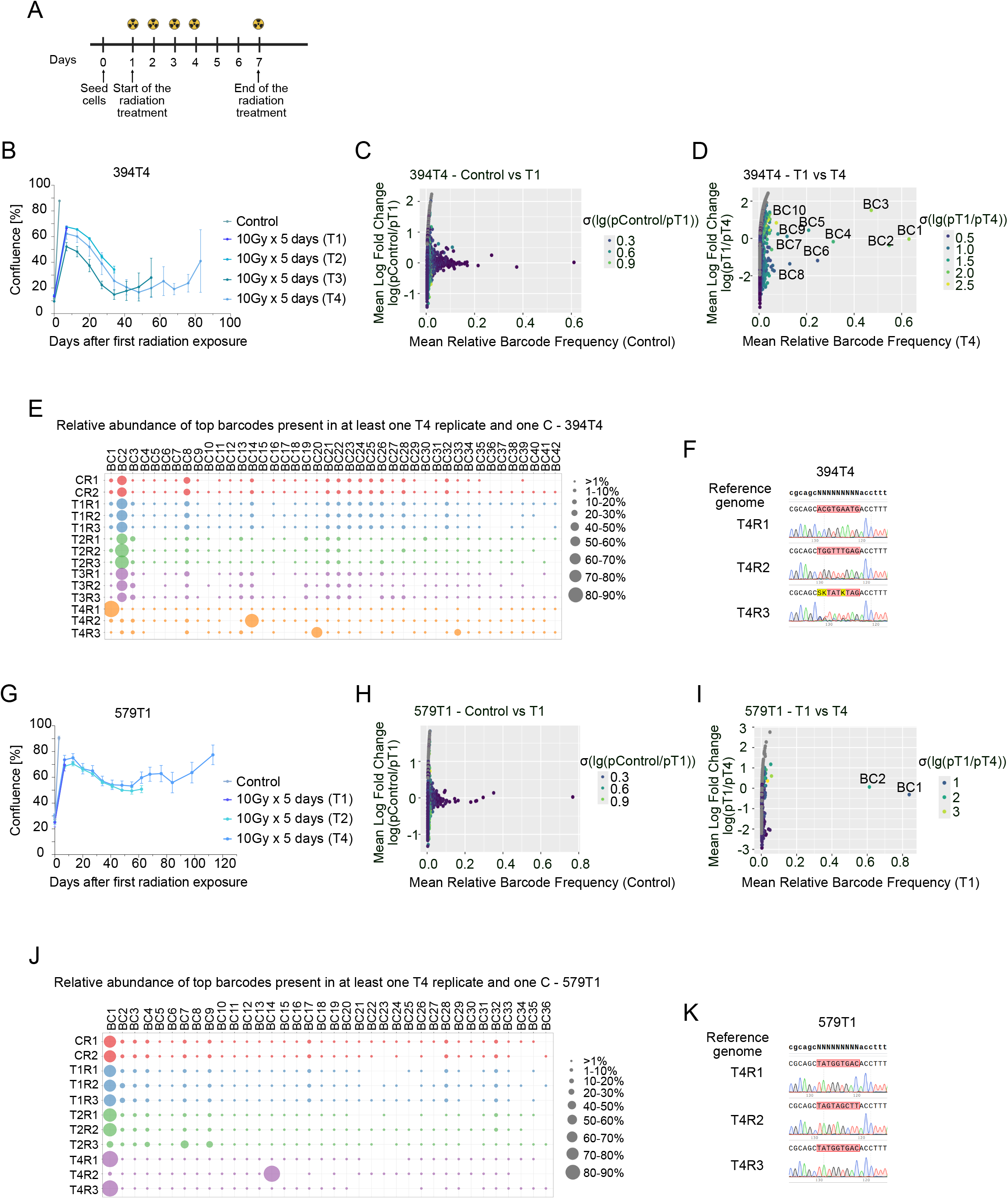
Radiation survival is a clonal event that reveals two distinct clonal trajectories. **(A)** Experimental timeline of the fractionated irradiation regime (10 Gy 5 times over the course of 7 days). **(B, G)** Proliferation curves, measured as percentage of confluence, for the cell lines **(B)** 394T4, and **(G)** 579T1 under radiation as indicated. Color coding distinguishes the non-irradiated control from the populations harvested at different time points (T1-T4). Data points represent mean values ± SEM. **(C, D, H, I)** Scatter plots of relative barcode frequencies between time points. (**C,D**) 394T4: Control vs. T1 (**C**), T1 vs. T4 (**D**). (**H,I)** 579T1: Control vs. T1 (**H**), T1 vs. T4 (**I**). Log₂ fold change is plotted against mean barcode abundance. **(E, J)** Bubble plots depicting the relative abundance of the top barcodes present in at least one T4 replicate and one control replicate for **(E)** 394T4 and **(J)** 579T1. Color coding distinguishes the different time points, while bubble size corresponds to relative abundance (%). **(F, K)** Sanger chromatograms of the three independent replicates (n=3) aligned to the reference genome for **(F)** 394T4 and **(K)** 579T1. Within the replicates, the barcode region is highlighted in red to denote a complete match, whereas mismatches are indicated in yellow. In the reference genome, the barcode region is represented as NNNNNNNNN.

Following irradiation, almost all cell lines exhibited a marked reduction in viability, with confluency dropping below 10%, indicating cell death. Nevertheless, approximately 80 days after initiation of radiation exposure, most lines began to recover and reached confluency levels above 30% (**Figure 2B, G; Supplementary Figure S2A–D**). Two cell lines displayed distinct responses: 579T1 maintained relatively stable confluency throughout the experiment, indicating quiescence rather than cell death (**Figure 2G**), whereas 238N1 failed to recover, confirming its high radiosensitivity (**Supplementary Figure S2B**). These responses were consistent with their previously observed radiosensitivity, as 579T1 displayed the highest radioresistance, whereas 238N1 was the most radiosensitive line among those tested (**Figure 1A**).

To investigate clonal dynamics, we performed quantitative DNA sequencing of relative barcode abundances in independent replicates (n=2–3 per time point per cell line) at four time points: end of treatment (T1), when confluency dropped below 30% (T2), at the lowest confluency point (T3), and after recovery began (T4), alongside a non-irradiated control (**Figure 1B**).

Across all barcoded lines, no major differences in barcode frequencies were observed between the non-irradiated control (Control) and T1 (**Figure 2C, H; Supplementary Figure S2E, I**). Barcode representation remained stable during the phase of cell death or quiescence (T3) (**Supplementary Figure S2M–W**). However, this scenario markedly changed at T4, corresponding to the recovery phase (T1 vs. T4, **Figure 2D, I; Supplementary Figure S2F, J).** Here, analysis of barcode dynamics revealed two distinct patterns of enrichment. In 394T4 and 482N1 (**Figure 2D; Supplementary Figure S2F, O, R**), the identity of the enriched barcodes at T4 varied considerably between replicates, indicating highly variable trajectories. This was further supported by analysis of the 20 most frequent barcodes present in at least one control replicate (**Figure 2E; Supplementary Figure S2G**) and by Sanger sequencing of the recovering population (**Figure 2F; Supplementary Figure S2H**). At endpoint, a single barcode dominated across all replicates, but its identity differed between replicates, indicating variable trajectories and supporting a stochastic component to recovery.

Interestingly, in 394T4, one barcode (BC2) remained dominant through early time points (Control, T1, T2, T3) but lost dominance at T4, suggesting that the dominant clone during early recovery was not the one driving long-term survival. In contrast, 579T1 and 579DLN exhibited reproducible barcode enrichment. In 579T1, each replicate was dominated by a single barcode at T4, with the same barcode dominant in two of three replicates (**Figure 2J**). This was consistent with the stable confluency observed during the experiment, indicating persistence of a limited number of cells rather than a marked population decline. This was coherent with the proliferation curve, in which 579T1 maintained relatively stable confluency throughout the experiment, indicating the persistence of cells with limited proliferation rather than the marked population decline observed in the other cell lines. Nevertheless, the emergence of a single dominant barcode in each replicate indicates that, even in a model with relatively high survival, fractionated irradiation imposes strong selective pressure that can ultimately result in the dominance of individual barcodes. Sanger sequencing confirmed the presence of a single dominant barcode in all replicates (**Figure 2K**). 579DLN also displayed a highly reproducible trajectory, with co-dominance of two barcodes in both replicates, each accounting for ∼50% of the population at endpoint (**Supplementary Figure S2K, L**). Neither barcode was predominant at baseline, indicating selection occurred during recovery. The reproducible emergence of the same two barcodes across replicates indicates highly consistent selection in 579DLN.

Overall, these results demonstrate that fractionated irradiation acts as a potent, selective bottleneck that reshapes clonal architecture toward one or two dominant barcodes. Importantly, the trajectories leading to survival can follow one of two distinct paths: either a highly variable pattern of enrichment, consistent with a stochastic component to recovery, or the reproducible enrichment of specific pre-existing barcodes.

### Radiation survivors exhibit convergent functional impairments despite molecular heterogeneity

Next, we characterized the surviving populations (Rx) to evaluate whether clonal selection or adaptation impacts continued radiosensitivity and, most importantly, metastatic ability. Interestingly, all clonal Rx cell lines displayed higher sensitivity to radiation re-exposure across the evaluated dose range compared to their parental counterparts **(Figure 3A, B; Supplementary Figure S3A–C**). This sensitivity is particularly evident in the context of the barcode-tracing data: the shift toward single-lineage homogeneity was not driven by the positive selection of a classic radioresistant subpopulation. On the contrary, the ultimately dominant clone does not appear to harbor an inherently radioresistant phenotype, reinforcing the notion that fractionated irradiation exerts a highly convergent selective pressure. These findings support that surviving a severe radiation regimen does not confer a long-term fitness advantage regarding radioresistance. Instead, this evolutionary pressure results in a recovered population that is more radiosensitive than its parental, radiation-naïve counterpart.

**Figure 3.**
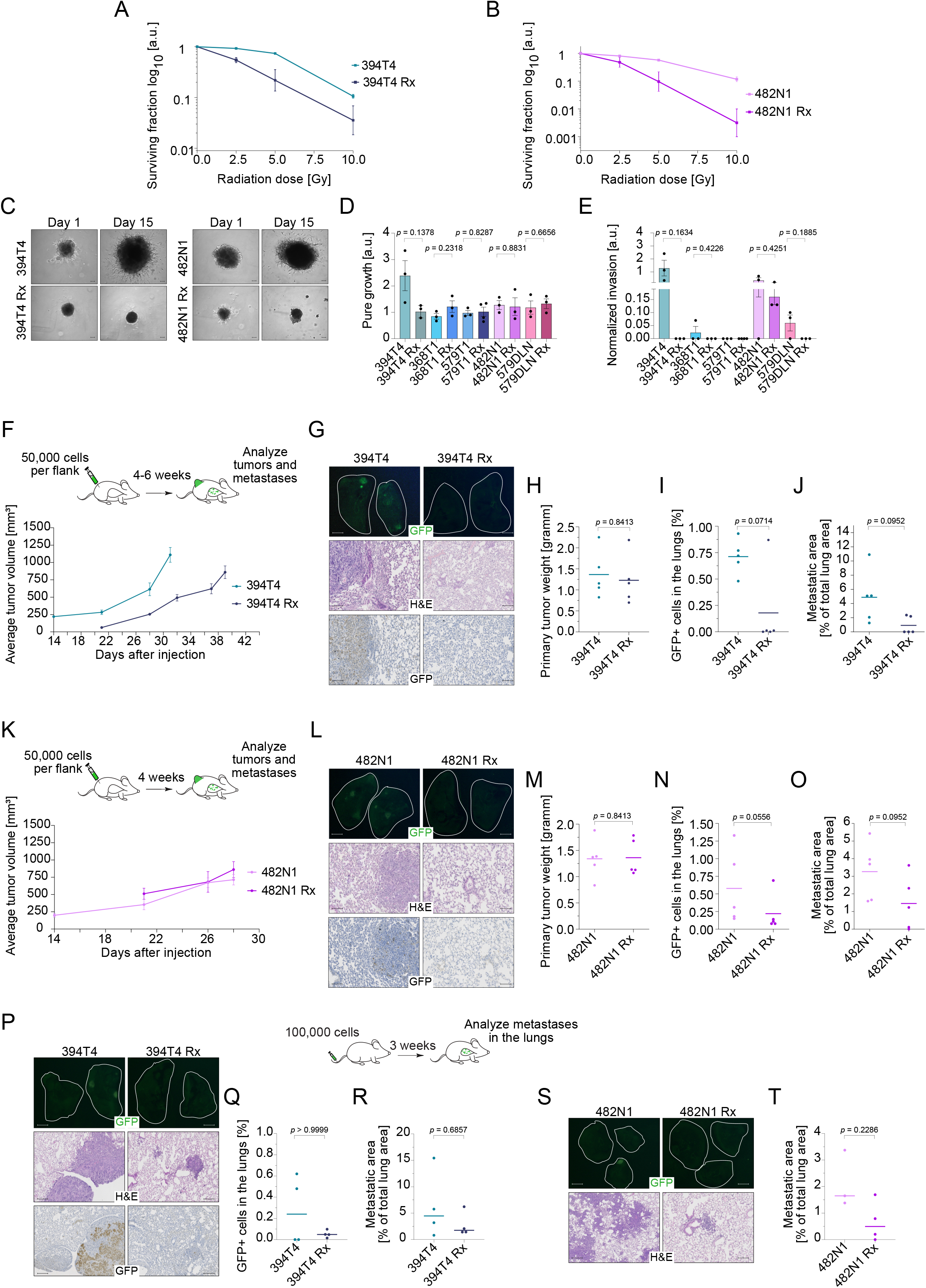
Radiation survivors exhibit increased radiosensitivity and impaired metastatic ability. **(A, B)** Radiation-induced clonogenic survival curves of parental versus radiation-surviving (Rx) counterparts for **(A)** 394T4 and **(B)** 482N1 cell lines. Data represent Mean ± SEM from N=3 independent biological replicates. **(C–E)** Spheroid invasion assay in Matrigel^®^. **(C)** Representative brightfield images at days 1 and 15 post-embedding (Scale bar: 200 µm). **(D)** Quantification of spheroid growth and **(E)** normalized invasion area. Data represent Mean ± SEM from N=3 independent biological replicates. *p*-values were determined using an unpaired Student’s t-test with Welch’s correction. **(F–O)** Subcutaneous model assessing primary tumor growth and lung metastasis in 394T4 (**F–J**) and 482N1 (**K–O**). (**F, K**) Primary tumor growth curves over time. (**G, L**) Representative fluorescence images of lungs showing GFP⁺ metastases (top), H&E (middle), and anti-GFP IHC (bottom) sections. Scale bars: 100 µm. (**H, M**) Tumor weight at endpoint. (**I, N)** Flow cytometry quantification of GFP⁺ cells in lungs. (**J, O**) Histological quantification of metastatic area relative to total lung area. **(P–T)** Intravenous tail-vein injection model for parental and radiation-surviving (Rx) 394T4 **(P–R)** and 482N1 **(S, T)** cells. **(P, S)** Representative fluorescence images of GFP (top) and H&E-stained lung sections (bottom). Scale bars: 100 µm. **(Q)** Quantification of GFP^+^ cells in lungs by flow cytometry. **(R, T)** Histological scoring of total metastatic burden. All dot plots display individual biological replicates (N=3-5), horizontal lines represent Mean ± SEM. *p*-values were determined by two-tailed Mann-Whitney U test.

We then evaluated whether radiation treatment had impacted the metastatic potential of the surviving populations. *In vitro*, we assessed spheroid growth and invasion using a 3D matrix-embedded spheroid assay (**Figure 3C–E; Supplementary Figure S3D–F**). Across all lines, Rx cells displayed no meaningful differences in spheroid core expansion compared with parental cells, with the exception of 394T4, where parental spheroids continued to grow while Rx spheroids remained essentially static. All other lines showed comparable growth between parental and Rx conditions (**Figure 3D**). In contrast, invasive behavior was uniformly lost: every cell line that initially generated invasive spheroids in the parental state failed to invade after radiation selection, indicating a consistent and robust loss of invasive capacity across all Rx populations (**Figure 3E**).

To determine whether radiation-survivor (Rx) populations retained metastatic competence *in vivo*, we evaluated metastatic ability using two complementary syngeneic immunocompetent models. Subcutaneous injection allowed assessment of the complete metastatic cascade, from escape from an established primary tumor to colonization and outgrowth at distant sites, while intravenous injection isolated the lung-colonization step. To avoid immune rejection due to sex mismatch, we confirmed the sex of origin for each cell line using *Rbm31x/y* PCR and injected cells only into mice of the corresponding sex (**Supplementary Figure S3G**).

Comparing parental and Rx 394T4 cells revealed a clear shift in primary tumor dynamics. Rx tumors showed a one-week delayed onset and altered growth kinetics (**Figure 3F**), but ultimately reached comparable endpoint weights (**Figure 3H**). This indicates that radiation survival is associated with impaired early engraftment and altered tumor growth dynamics, without preventing the eventual formation of tumors of comparable size. Despite this preserved growth, systemic dissemination was strongly reduced. Lung metastasis was abundant in parental tumors but nearly absent in the Rx group (**Figure 3G, I, J),** as confirmed by GFP-FACS quantification and histological analyses. The metastatic 482N1 line showed the same overall pattern: delayed Rx tumor onset and altered growth kinetics, followed by comparable endpoint tumor weights and a marked reduction in lung metastatic burden (**Figure 3K–O**).

To test whether this impairment persisted when bypassing early metastatic steps, we performed intravenous injections. Across all lines, Rx populations showed a drastic loss of lung colonization, frequently approaching complete absence of metastases (**Figure 3P–T; Supplementary Figure S3H–R**). Non-barcoded 394T4 and 368T1 parental and Rx controls confirmed that this phenotype was not barcode-dependent (**Supplementary Figure S3H, I, Q, R**).

Together, these results demonstrate that survival after fractionated irradiation is not associated with increased radioresistance but instead leads to a consistent loss of invasive and metastatic fitness. Rx cells display a transient delay in primary tumor establishment yet ultimately form tumors of similar final burden, while failing to execute the metastatic cascade, even when directly delivered to the lungs, indicating a generalized post-irradiation collapse of metastatic competence across all NSCLC models.

### Transcriptional and metabolic recovery occurs through distinct, lineage-specific programs

The functional assays revealed a shared phenotype across NSCLC radiation-survivor (Rx) populations: increased sensitivity to re-irradiation and loss of invasive and metastatic capacity. We next asked whether this convergence extended to the transcriptional level or whether distinct molecular routes underlay radiation survival.

To profile acute and recovery responses, we performed bulk RNA-sequencing at control, T1 (immediately post-irradiation), and T4 (recovery) time points (**Figure 4, Supplementary Figure S4**). Unsupervised hierarchical clustering of the top 500 highly variable genes revealed pronounced cell-line- and time-specific transcriptional dynamics (**Figure 4A, D; Supplementary Figure S4I, L, W, Di**). 394T4 showed minimal acute remodeling, whereas 482N1 underwent a strong transcriptional shift at T1. All lines ultimately established T4 states clearly distinct from their parental controls, but the trajectories leading to these states differed substantially.

**Figure 4.**
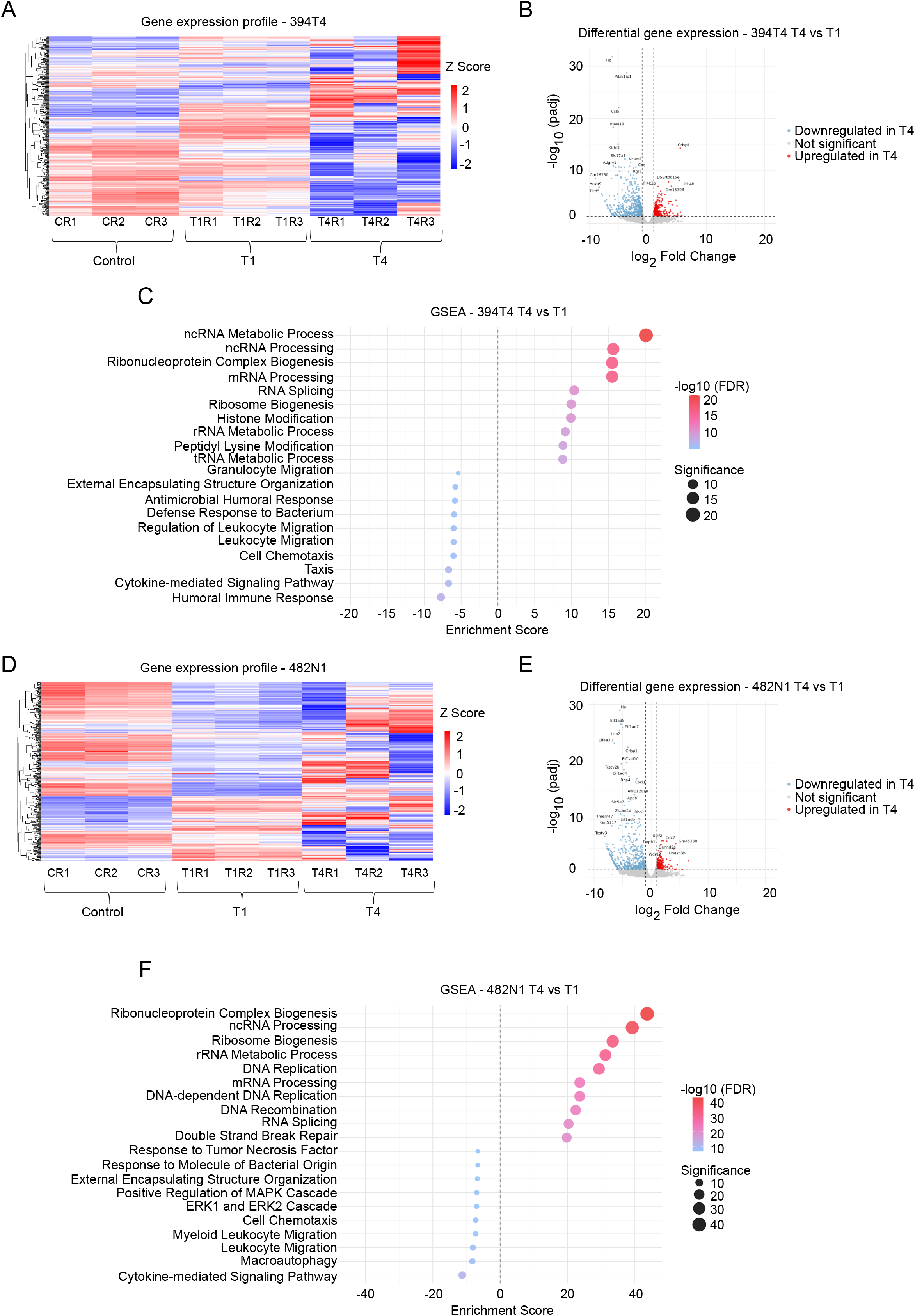
Radiation survivors converge on proliferative recovery through divergent transcriptional trajectories. **(A, D)** Hierarchical unsupervised clustering heatmaps of the top 500 highly variable genes for (**A**) 394T4 and (**D**) 482N1. Genes were pre-filtered (mean expression < 10) and selected based on highest row variance. Samples include baseline (Control; CR1–CR3), acute (T1; T1R1–T1R3), and recovery (T4; T4R1–T4R3) phases. "R" denotes independent biological replicates. Color intensity represents relative gene expression levels scaled as Z-scores, shifting from downregulated (blue) to upregulated (red) states. **(B, E)** Volcano plots illustrate differentially expressed genes (DEGs) in the recovery phase (T4) relative to the acute phase (T1) for **(B)** 394T4 and **(E)** 482N1 cell lines. Dashed vertical and horizontal lines indicate the log_2_ fold-change > 1 and statistical significance thresholds (*p*-adjusted < 0.05), respectively. Statistically significant upregulated genes are highlighted in red, whereas downregulated genes are shown in blue. **(C, F)** Gene Set Enrichment Analysis (GSEA) bubble plots displaying significantly enriched biological processes in the recovery phase (T4) relative to the acute phase (T1) for **(C)** 394T4 and **(F)** 482N1 cell lines. The x-axis indicates the normalized enrichment score (NES) for each gene set, where positive and negative values represent enrichment and depletion, respectively. Both bubble size and color intensity reflect the statistical significance scale, quantified as −log_10_ (FDR), shifting from low significance (blue) to high significance (red).

Importantly, transcriptional heterogeneity at T4 did not simply mirror barcode composition. Lines dominated by a single barcode (e.g., 579T1) still displayed substantial transcriptional variability, whereas others with multiple dominant barcodes (e.g., 579DLN) remained comparatively homogeneous. Thus, although fractionated irradiation imposed a strong bottleneck that reshaped barcode composition, the resulting barcode distribution did not show a simple correspondence with the transcriptional state of the recovered populations. This suggests that barcode composition alone was insufficient to explain the transcriptional heterogeneity observed.

We next analyzed differential gene expression and pathway enrichment to define the biological processes underlying these trajectories. Despite cell-line-specific differences, the acute response showed notable pathway-level convergence. Across NSCLC, GSEA revealed induction of chromosome segregation, organelle fission, and mitotic programs at T1, together with repression of ribosome biogenesis and RNA-processing pathways (**Supplementary Figure S4B, F, P, T, Ai, Fi).** These signatures were consistent with radiation-induced cell-cycle disruption rather than effective proliferation. Individual lines displayed additional stress- or inflammation-associated programs: antioxidant and chaperone induction in 394T4, inflammatory and injury-related genes in 482N1, ribosomal repression in 579T1, and a distinct stress signature in the non-recovering 238N1 (**Supplementary Figure S4A, E, O, S, Ei**). In contrast, 368T1 showed minimal acute transcriptional remodeling (**Supplementary Figure S4Z**). Thus, a partially shared cell-cycle–associated response was embedded within diverse stress, inflammatory, and metabolic programs.

During recovery, all surviving populations converged on transcriptional signatures associated with proliferative reactivation, including enrichment of organelle fission, chromosome segregation, and mitotic pathways, accompanied by downregulation of ribosome biogenesis and cytoplasmic translation (**Supplementary Figure S4C, D, G, H, Q, R, U, V, Bi, Ci**). Although several of these pathways were also enriched at T1, their biological context differed: at T1 they accompanied declining confluency, whereas at T4 they coincided with resumed growth.

Longitudinal comparisons revealed that the molecular routes to recovery differed between lines. In 394T4, T4 was marked by enrichment of RNA-processing and ribosome-biogenesis pathways and resolution of cytokine-associated signaling (**Figure 4B, C**). In 482N1, recovery involved ribosome biogenesis and DNA replication together with attenuation of acute inflammatory and ERK1/2-associated programs (**Figure 4E, F**). Other lines displayed distinct metabolic and mitochondrial signatures, ranging from reduced OXPHOS in 368T1 to increased oxidative phosphorylation and fatty-acid metabolism in 579T1, and aerobic-respiration enrichment in 579DLN (**Supplementary Figure S4Y, K, N**).

To functionally validate these transcriptional changes, we profiled redox balance, mitochondrial activity, and cellular energy status. Most Rx lines showed elevated cytosolic ROS (**Figure 5A–E**), indicating chronic oxidative stress, although 579T1 did not, suggesting cell-line-specific redox adaptation. Mitochondrial ROS and membrane potential varied markedly between models (**Figure 5F–O**). Some lines combined increased mtROS with mitochondrial hyperpolarization (394T4, 368T1), others showed reduced mtROS with depolarization (482N1), and still others displayed high membrane potential despite low mtROS (579T1, 579DLN). Thus, mitochondrial polarization, oxidative stress, and ROS production were uncoupled across survivors.

**Figure 5.**
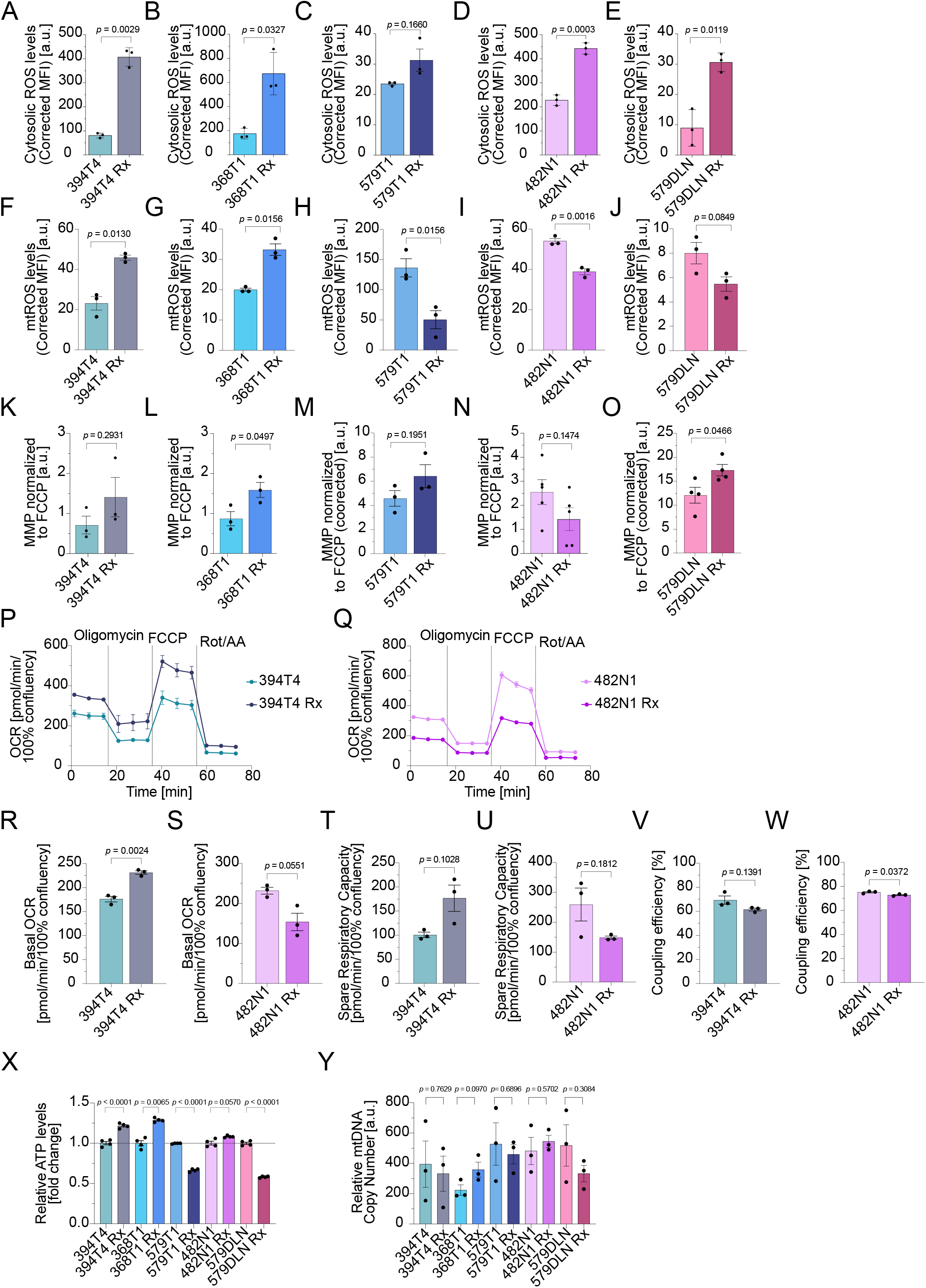
Radiation survival is associated with distinct bioenergetic states. **(A-E)** Quantitative cytosolic reactive oxygen species (ROS) levels evaluated via flow cytometry using CellROX Orange/Deep Red or CM-H2DCFDA for **(A)** 394T4, **(B)** 368T1, **(C)** 579T1, **(D)** 482N1, and **(E)** 579DLN cell lines. **(F-J)** Quantitative mtROS levels evaluated with flow cytometry using MitoSOX Red or MitoSOX Green for **(F)** 394T4, **(G)** 368T1, **(H)** 579T1, **(I)** 482N1, and **(J)** 579DLN cell lines. For both ROS and mtROS assays, the y-axis displays background-corrected Median Fluorescence Intensity (MFI) expressed in arbitrary units [a.u.]. Data represent Mean ± SEM from n=3 technical replicates within one out of three biological replicates (N=3). **(K-O)** Quantitative mitochondrial membrane potential (MMP) evaluated via flow cytometry using TMRM for **(K)** 394T4, **(L)** 368T1, **(M)** 579T1, **(N)** 482N1, and **(O)** 579DLN cell lines. The y-axis displays background-corrected MFI expressed in arbitrary units [a.u.] and normalized to the FCCP-uncoupled control. Data represent Mean ± SEM from N=3-5 independent biological replicates. **(P-W)** Mitochondrial respiration profiles using a Seahorse Xfe96 analyzer for 394T4 and 482N1 cell lines. **(P, Q)** Oxygen consumption rate (OCR) kinetics, normalized to cell confluence, measured over time under baseline conditions and following successive injections of Oligomycin, FCCP, and Rotenone/Antimycin A (Rot/AA) for **(P)** 394T4, and **(Q)** 482N1. **(R, S)** Basal respiration rates, **(T, U)** spare respiratory capacity, **(V, W)** coupling efficiency (%) for **(R, T, V)** 394T4 and **(S, U, W)** 482N1. Seahorse kinetic data represent n=4 technical replicates within one out of three biological replicates, bar graphs represent Mean ± SEM from N=3 independent biological replicates. **(X)** Relative intracellular ATP levels across the indicated NSCLC cell lines, expressed as fold-change relative to their respective non-irradiated parental counterparts. **(Y)** Relative mtDNA copy number determined via quantitative real-time PCR (qPCR) and expressed in arbitrary units [a.u.]. Data represent Mean ± SEM from N=3 independent biological replicates. *p*-values were determined using an unpaired Student’s t-test with Welch’s correction.

Seahorse analysis further confirmed divergent respiratory states (**Figure 5P–W; Supplementary Figure S5A–L**). Some Rx populations maintained higher basal respiration (394T4, 368T1), whereas others showed reduced oxygen consumption (579T1, 482N1, 579DLN). Spare respiratory capacity also differed, with only 394T4 Rx showing an increased reserve. Despite these differences, coupling efficiency remained largely unchanged.

These mitochondrial states translated into distinct energetic outputs. ATP levels increased in 394T4 and 368T1 but decreased in 579T1 and 579DLN, while 482N1 maintained parental ATP levels despite reduced respiration (**Figure 5X**). mtDNA copy number remained unchanged in all cell lines (**Figure 5Y**), indicating that metabolic rewiring reflected functional rather than quantitative mitochondrial changes.

Overall, metabolic profiling revealed three bioenergetic states among NSCLC radiation survivors: 1) high-output mitochondria with increased respiration and ATP (394T4, 368T1), 2) low-output mitochondria with reduced respiration and ATP but preserved polarization (579T1, 579DLN), and 3) an intermediate state maintaining ATP despite reduced mitochondrial activity (482N1). Thus, radiation survival does not produce a uniform metabolic phenotype but instead drives divergent mitochondrial and energetic configurations that support long-term persistence through distinct metabolic strategies.

To determine whether the metabolic states of radiation survivors were reflected at the transcriptional level, we performed unsupervised hierarchical clustering of the top 500 T4 genes (**Figure 6A**). 579T1 and 579DLN cells clustered closely together, whereas 394T4, 368T1, and 482N1 grouped within a separate branch, with 482N1 branching separately from 394T4 and 368T1. This transcriptional organization was consistent with the three bioenergetic states identified by our functional metabolic assays.

**Figure 6.**
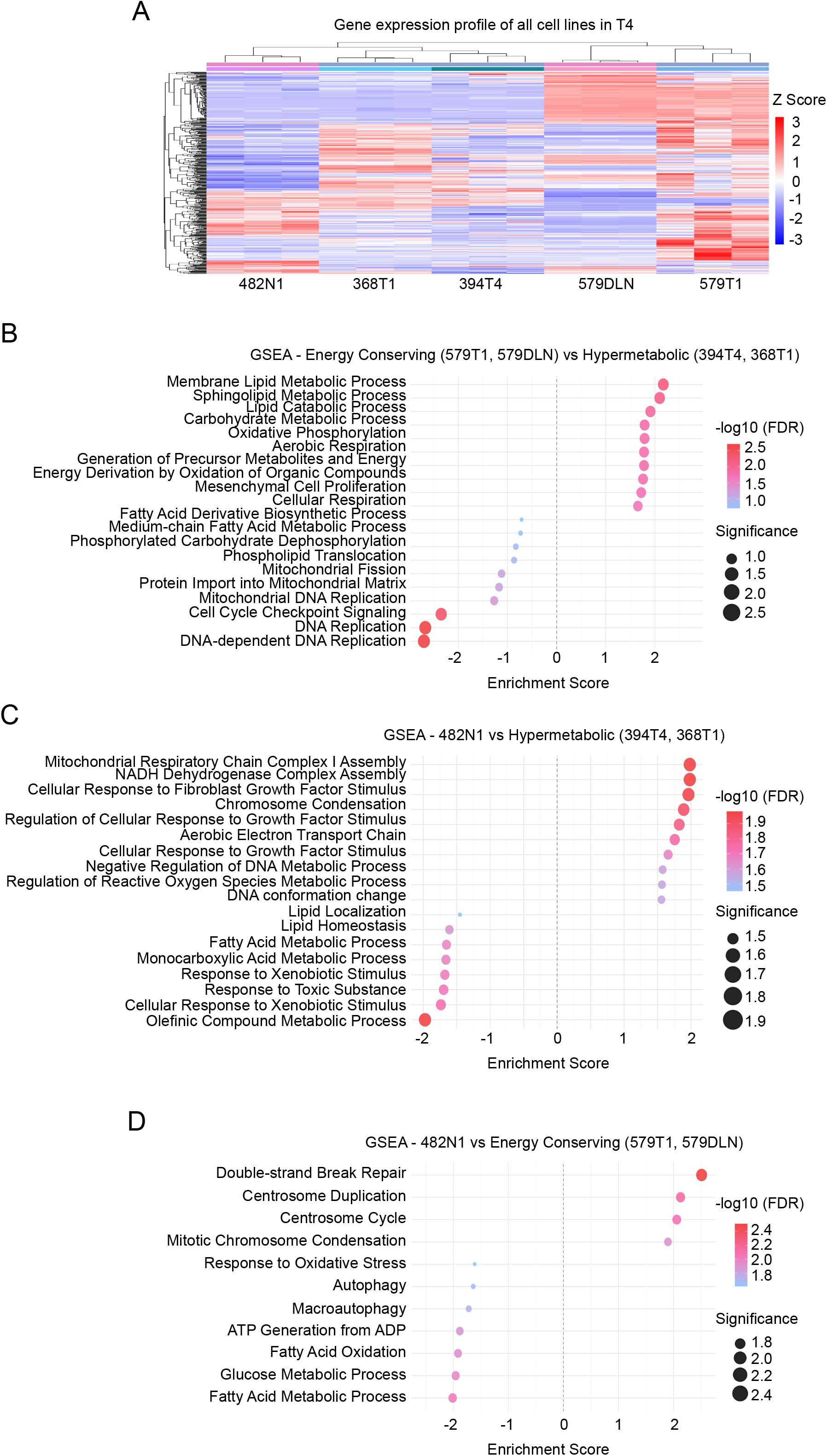
Radiation survival is associated with distinct bioenergetic states. **(A)** Hierarchical clustering heatmap displaying the top 500 highly variable genes across all evaluated cell lines (482N1, 368T1, 394T4, 579DLN, and 579T1) during the recovery phase (T4). Both rows (genes) and columns (independent biological replicates) were grouped using unsupervised hierarchical clustering to reveal overarching expression patterns. Color intensity represents relative gene expression levels scaled as Z-scores, shifting from downregulated (blue) to upregulated (red) states. **(B-D)** Gene Set Enrichment Analysis (GSEA) displaying significantly enriched biological processes for distinct phenotypic comparisons. **(B)** Energy-conserving cell lines (579T1, 579DLN) compared against Hypermetabolic cell lines (394T4, 368T1). **(C)** 482N1 compared against Hypermetabolic cell lines (394T4, 368T1). **(D)** 482N1 compared against Energy-conserving cell lines (579T1, 579DLN). For all GSEA plots, the x-axis indicates the normalized enrichment score (NES) for each gene set, where positive and negative values represent enrichment and depletion, respectively. Both bubble size and color intensity reflect the statistical significance scale, quantified as −log_10_ (FDR), shifting from low significance (blue) to high significance (red).

We next compared transcriptional programs between the hypermetabolic (394T4, 368T1), energy-conserving (579T1, 579DLN), and metabolically suppressed (482N1) groups. Energy-conserving cell lines showed enrichment of lipid metabolic and mitochondrial maintenance pathways, including increased expression of *Rab5if*, *ND3*, and *Mrpl34* (**Figure 6B; Supplementary Figure S6A**). In contrast, hypermetabolic cell lines upregulated DNA-replication and proliferation-associated pathways and expressed oxidative-stress and remodeling genes such as *SOD1* and *Septin2*. Comparisons with 482N1 revealed a distinct third transcriptional state. Relative to hypermetabolic cell lines, 482N1 was enriched for mitochondrial-complex remodeling, ROS-regulatory pathways, and developmental regulators (*Pax2*, *Six1*, *Eomes*, *Fgfr1*) (**Figure 6C; Supplementary Figure S6B).** Relative to energy-conserving cell lines, 482N1 showed strong enrichment of mitotic, meiotic, and DNA-repair programs, whereas the energy-conserving phenotype remained dominated by metabolic and mitochondrial-homeostasis pathways (**Figure 6D; Supplementary Figure S6C**).

Together, these analyses reveal that although multiple radiation-surviving cell lines share reduced mitochondrial activity, their transcriptional solutions differ. Energy-conserving cell lines maintain lipid and mitochondrial homeostasis programs, whereas 482N1 adopts a regulatory and cell-cycle–associated transcriptional state while remaining metabolically suppressed. Thus, radiation survival produces three distinct transcriptional–metabolic phenotypes, reflecting divergent molecular trajectories despite shared functional constraints.

### Functional convergence is not driven by clonal dominance

Crucially, the transcriptional and metabolic states of radiation-surviving populations were not determined by clonal dominance. Despite the emergence of a single dominant barcode in 579T1, this line clustered separately from 394T4 - a line with multiple dominant barcodes - in unsupervised hierarchical clustering (**Figure 6A**). Similarly, 482N1, which lacked a dominant barcode at endpoint, exhibited a distinct transcriptional signature enriched in mitotic and DNA repair programs (**Figure 6C, D**). These findings demonstrate that the molecular and functional phenotype of radiation survivors is not dictated by clonal composition alone. Rather, it arises from a dynamic interplay between clonal selection and post-irradiation reprogramming, a process that operates independently of lineage dominance. This decoupling explains how diverse clonal trajectories can converge on shared functional outcomes: increased radiosensitivity and impaired metastatic fitness, the very hallmarks of the radiation-surviving state.

## DISCUSSION

Radiation response is not uniform across tumor cell populations, reflecting the inherent heterogeneity of cancer and its capacity to withstand treatment-induced stress (11). In our NSCLC model, cell lines displayed markedly different radiosensitivities despite sharing a common genetic background. This heterogeneity is clinically relevant, as fractionated radiotherapy imposes repeated selective pressure that may favor distinct survival strategies (11–13). The regimen used, 5 × 10 Gy over 7 days, mimics clinically relevant hypofractionated schedules, such as stereotactic body radiotherapy (SBRT) (14, 15), and was chosen to induce a pronounced clonal bottleneck without complete eradication.

Using longitudinal cellular barcoding, we found that fractionated irradiation imposed a strong clonal bottleneck, but the trajectories to recovery followed two distinct patterns: stochastic enrichment in some lines (e.g., 394T4, 482N1) and reproducible selection of pre-existing barcodes in others (e.g., 579T1, 579DLN). Notably, the dominant barcode at endpoint was not always the most abundant at baseline, indicating that survival was not solely determined by initial clonal fitness. These findings align with previous studies showing that radioresistance can arise through both stochastic and deterministic mechanisms (16), but they reveal a critical new layer: **clonal selection is not predictive of functional outcome**.

However, these distinct clonal dynamics did not result in distinct phenotypes. Following recovery from fractionated irradiation, all evaluated Rx populations were more sensitive to subsequent irradiation than their parental counterparts. This indicates that survival to the radiation treatment did not confer an advantage upon re-exposure. Previous studies have reported that radiosensitivity and metastatic potential are not necessarily linked (17), and that fractionated irradiation does not necessarily select for increased radioresistance or metastatic activity (18). Our study extends these observations by directly characterizing the functional properties of populations that survived and recovered from this treatment. Despite the strong clonal selection observed, the surviving populations showed reduced invasive capacity and metastatic fitness. Primary tumor establishment was delayed, although tumors eventually reached comparable burdens, while lung metastatic colonization was markedly reduced. Importantly, this impairment persisted following intravenous injection, indicating that it was not restricted to the early steps of the metastatic cascade. These findings argue against a direct association between radiation survival and metastatic capacity, and indicate that radiation survival did not confer a broader fitness advantage, but was instead associated with increased sensitivity to subsequent radiation exposure and reduced invasive and metastatic potential.

Despite these common functional outcomes, the transcriptional responses of the surviving populations were only partially shared. During the acute response, enrichment of chromosome segregation, organelle fission, and mitotic pathways across cell lines was consistent with a common response to radiation-induced cell-cycle disruption rather than active proliferation, in line with the activation of DNA damage checkpoints and transient cell-cycle arrest following ionizing radiation (19, 20).

Metabolic profiling confirmed three distinct bioenergetic states among survivors: hypermetabolic (394T4, 368T1), energy-conserving (579T1, 579DLN), and metabolically suppressed (482N1). These states were supported by unique transcriptional programs and were not predicted by barcode dominance. Notably, mtDNA copy number remained unchanged, indicating that metabolic rewiring reflects functional adaptation rather than quantitative mitochondrial changes (6, 21–24). Moreover, these metabolic and redox responses can vary depending on the cellular context (25, 26). Such metabolic changes may contribute to cellular adaptation and survival under oxidative stress (27). Thus, the transcriptomic analyses supported the distinct phenotypes identified by metabolic assays. Together, these findings indicate that radiation survival is associated with different bioenergetic configurations, suggesting that surviving populations can adapt to the metabolic demands of recovery through distinct strategies. Despite these differences, a common feature of the surviving populations is reduced overall metabolic plasticity, which allows cancer cells to adapt their metabolism to changing energetic demands during tumor progression and metastasis (28–30). Therefore, reduced metabolic flexibility might explain the common phenotype of reduced metastatic ability observed in radiation survivors, as this complex process requires high metabolic flexibility to adapt to changing energetic demands.

These results have important implications for therapy. Rather than assuming that radiation selects for resistant clones, we should consider that it may suppress metastatic spread. Future strategies could exploit this vulnerability — for example, by combining radiation with agents that target the metabolic or transcriptional states of survivors to prevent relapse. Moreover, the decoupling of clonal selection and adaptation suggests that targeting adaptive pathways — not just resistant clones — may be key to improving outcomes.

One limitation is that metabolic states were inferred from transcriptional and functional assays, not direct metabolite or flux measurements. Future work should employ metabolomics and flux analysis to define the metabolic pathways supporting each state. Additionally, functional validation of key transcriptional regulators and the long-term stability of the post-irradiation phenotype remain open questions.

In summary, radiation survival in NSCLC does not generate a uniform resistant phenotype. Instead, it triggers a clonal bottleneck that leads to diverse molecular and metabolic trajectories, converging only on functional outcomes that may be therapeutically exploitable.

## METHODS

### Data availability statement

All information and requests for resources and reagents should be directed to the lead contact, Barbara M. Grüner. All sequencing data (RNA-seq from cell lines under radiation therapy, and DNA-Barcode sequencing) is available on NCBI (Bioproject number: PRJNA1492111).

### Cell lines

The 368T1, 394T4, and 579T1 cell lines were generated from primary lung tumors, 238N1 and 482N1 from lymph node metastases, and 579DLN from a distant lymph node metastasis of *Kras^LSL-G12D/+^;Trp53^flox/flox^* (*Kras^G12D^;Trp53^KO^*) *Kras^LSL-G12D+^;Trp53^flox/flox^;Rosa26^LSL-Tomato/+^* (*Kras^G12D^;Trp53^KO^;R26^Tomato^*) genetically engineered mice (9, 31). Notably, 579T1 and 579DLN were derived from the same mouse.

Human embryonic kidney (HEK) 293T cells (ATCC, Cat# CRL-11268) were utilized as packaging cells for lentiviral production. All cell lines were regularly tested and confirmed to be *Mycoplasma* negative, and they were passaged at least once after thawing before their use in subsequent experiments. Cell lines were maintained in culture no longer than 1 month. After this period, a new vial of cells was thawed. All cells were cultured in Dulbecco’s Modified Eagle Medium (DMEM; Gibco) supplemented with 10% Fetal Bovine Serum (FBS; VWR/Biowest), 50 U/ml penicillin, 50 µg/ml streptomycin, and 1 mM L-glutamine (Thermo Fisher Scientific).

### X-Ray Irradiation

Irradiation was performed at room temperature using an RS320 cabinet irradiator (Xstrahl, Ltd., UK, RRID:SCR_024344) at a source-to-surface distance (SSD) of 50 cm. The system operated at a tube voltage of 320 kV and a current of 10 mA with an unfiltered X-ray beam, delivering radiation at a constant dose rate of approximately 1.0 Gy/min.

### Colony formation assay

Colony Formation Assay (CFA), or clonogenic assay, was performed to determine the radiotolerance or radiosensitivity of the above-mentioned cell lines. Cells were counted (Countess II FL, Invitrogen, RRID:SCR_018591) and seeded at an optimized low density (100-400 cells per well) in 6-well plates, using three replicates per dose. At 24 hours post-seeding, cells were irradiated at doses of 2.5, 5 and 10 Gy. A non-irradiated plate was used as a control for downstream analysis. Following irradiation, cells were kept in the incubator at 37 °C and 5% CO_2_.

Approximately 7-10 days post-irradiation, when colonies reached a size of ∼50 cells and were visible in all conditions, the plates were fixed and stained with crystal violet (Sigma-Aldrich, Cat# HT90132) for colony counting. Plates were scanned and colonies quantified using the Cytation 5 Imaging Multi-Mode Reader (Agilent Technologies, RRID:SCR_019732) with Gen 5™ Microplate Reader and Imager Software (version 3.14, Agilent Technologies, RRID:SCR_017317) for automated counting. Cell size and intensity thresholds were adjusted for each cell line to ensure accurate automated quantification. The plating efficiency and surviving fraction (SF) were calculated as described elsewhere (10). All assays were performed in at least three independent biological replicates.

### Cloning of the sCORE (Single Cell Clone Tracer) library

The vector backbone, pLenti CMV GFP Hygro (Addgene plasmid # 17446; http://n2t.net/addgene:17446; RRID:Addgene_17446), was prepared by overnight digestion at 37 °C with PspOMI (New England Biolabs (NEB), Cat# R0565S) and Nhel-IHF (NEB, Cat# R3131S) and dephosphorylation with antarctic phosphatase (NEB, Cat# M0289S), followed by gel purification using a 2% agarose Gel and the QIAquick^®^ Gel Extraction Kit (QIAGEN, RRID:SCR_021966). The resulting DNA was eluted in 30 μl of H_2_O.

The DNA insert containing the lineage barcodes was generated by PCR amplifying a ssDNA Oligo containing a random 9-bp-long barcode sequence and the NotI and Nhel-IHF cutting sites, followed by gel purification using a 2% agarose Gel the QIAquick^®^ Gel Extraction Kit (QIAGEN, RRID:SCR_021966) and elution in 30 µL H_2_O. The resulting dsDNA was then digested overnight at 37 °C with NotI (NEB, Cat# R0189S) and Nhel-IHF (NEB) and purified using the same QIAquick^®^ Gel Extraction Kit.

To generate the sCORE library (**Figure 1B**) the barcode-containing DNA insert and the backbone were mixed in a 3:1 insert:vector ratio and ligated together overnight using temperature-cycle ligation alternating between 10 °C and 30 °C in 30-second cycles and the T4 DNA ligase (NEB, Cat# M0202S). The ligation product was purified using the QAIQuick PCR purification Kit (Qiagen, RRID:SCR_021967) and eluted in 20 µL H_2_O.

For the plasmid pool generation, chemically competent E.coli One Shot™ Stbl3™ (Invitrogen, Cat# C737303) were used for transformation. To ensure high barcode library complexity and distribution, 17 transformation reactions with 0.2-1 µg of introduced DNA each were pooled and incubated as liquid cultures with 100 µg/mL Ampicillin overnight for ∼16 hours, at 37°C and 180 rpm using an Ecotron incubator shaker (INFORS-HT, Cat# 69400). Plasmid isolation from the bacterial cells was performed using the QIAGEN Plasmid Plus Mega Kit (Qiagen, RRID:SCR_025218) following the manufacturer’s instructions. Sanger Sequencing (performed by Microsynth Seqlab) of a few colonies was used to validate the success of the cloning approach. Library complexity of the final plasmid pool was estimated using NGS sequencing (Illumina MiniSeq).

### Lentivirus production (HEK293T transfection)

Lentiviral particles were produced as described before (32) by transfecting human embryonic kidney (HEK) 293 T cells (RRID:CVCL_0063) with the pLenti CMV GFP Hygro 9N transfer vector as well as packaging and envelope plasmids (pMD2.G, Addgene plasmid #12259; pMDLg/pRRE, Addgene plasmid #12251; and pRSV-Rev, Addgene plasmid #12253) using the transfection reagent TransIL-LT1 (Mirus Bio, Cat# MIR 6600) in combination with Opti-MEM medium (Thermo Fisher Scientific, Cat# 31985062). A negative control (no transfer vector) was included to confirm the absence of background signal.

### Generation of barcoded cell lines

394T4, 482N1, 579T1, and 579DLN cells were transduced with lentiviral sCORE particles at a low Multiplicity of Infection (MOI) (between 0.1 and 0.2) to minimize the probability of multiple barcode integrations per cell. Transduction efficiency was evaluated by flow cytometry based on GFP expression, using a BD FACSCelesta^TM^ Flow Cytometer (BD Biosciences, RRID:SCR_019597). Subsequently, barcoded cells were enriched via fluorescence-activated cell sorting (FACS) using an S3^TM^ Cell Sorter (Bio-Rad Laboratories, RRID:SCR_020224) to remove the non-barcoded population. The sorted cells were subsequently expanded, and their GFP expression was re-evaluated using a BD FACSCelesta™ flow cytometer (BD Biosciences). Only cell populations with > 90% GFP-positive cells were used for downstream experiments. Representative plots and the flow cytometer gating strategy are shown in **Supplementary Figure S7A**.

### Longitudinal monitoring of cell populations under fractionated irradiation

Barcoded cells were seeded in 12-well plates (Sarstedt, Cat# 83.3921) at a density of ∼10-30% confluency 24 hours post-seeding. Irradiation was administered using a consistent dose of 10 Gy 5 times over the course of 7 days, and cell confluence was monitored throughout the experiment using a NYONE automated cell imager equipped with Synentec cell recognition software (Synentec GmbH, RRID:SCR_024623). Cell culture medium was changed once or twice weekly, contingent upon metabolic demand.

Cells were harvested in three independent replicates at four time points: immediately after the final irradiation (T1), when confluency dropped below 30% (T2), at the lowest confluency point (T3), and upon regrowth (T4). T1 corresponds to the end of the radiation regimen, before significant cell death. Additionally, non-irradiated cells were harvested at ∼80% confluency to serve as the control group (Control). Moreover, regrown cells were subsequently expanded for use in following assays and are collectively referred to hereafter as “radiation survivors” or “Rx” cells.

### Clonal dynamics analysis via DNA sequencing

Following the radiation and recovery experiments, genomic DNA (gDNA) was isolated from harvested cells from two to three independent replicates at indicated time points using the Monarch^®^ kit (New England Biolabs, RRID:SCR_025194) according to the manufacturer’s instructions. Isolated gDNA was quantified using a Qubit 3 Fluorometer (Thermo Fisher Scientific, RRID:SCR_020311) with the Qubit dsDNA High Sensitivity (HS) Assay Kit (Invitrogen, RRID:SCR_020235). For each sample and replicate, 200 ng of gDNA was amplified via polymerase chain reaction (PCR) using KAPA HiFi HotStart polymerase (Roche, Cat# 07958935001). Primers were designed to incorporate Illumina sequencing primer-binding sites, adapters, and multiplexing indexes, utilizing a universal forward primer and a sample-specific indexed reverse primer. PCR products were resolved via 2% agarose gel electrophoresis to confirm amplicon presence. The targeted 248 bp barcoded region was excised, and DNA was purified using the QIAquick Gel Extraction Kit (QIAGEN, RRID:SCR_021966). Purified amplicons were quantified via Qubit as described above, pooled in equimolar concentrations, combined with a 30% PhiX control (Illumina, Cat# FC-110-3001) spike-in, and sequenced on an Illumina MiniSeq System (Illumina, Inc., RRID:SCR_016379) utilizing the MiniSeq High Output reagent kit (300 cycles, paired-end; Illumina, Inc.).

Raw sequencing reads were processed using a custom Python pipeline (version 3.9.9, Python Software Foundation, RRID:SCR_008394) for adapter trimming, barcode extraction, and frequency quantification (33). Barcode counts were subsequently imported into RStudio (version 4.5.1, Posit, RRID:SCR_000432) for normalization and downstream analysis.

Differential abundance was calculated by comparing irradiated time points (T1 and T4) to non-irradiated controls, as well as through pairwise temporal comparisons to track the dynamics of clonal evolution. Clonal dominance shifts were evaluated by tracking the relative frequencies of the top 20 most abundant barcodes across replicates and experimental conditions.

### Sanger sequencing for barcode validation

To validate the clonal composition and evaluate clonal dominance at the experimental endpoint (T4), part of the same gDNA used for high-throughput sequencing was subjected to Sanger sequencing (Microsynth Seqlab). Chromatograms and sequencing reads were analyzed using SnapGene software (Dotmatics, RRID:SCR_015053).

### Transcriptomic analysis via RNA sequencing

To elucidate the molecular mechanisms underlying radiation survival, transcriptomic profiling was conducted on cells harvested from three independent replicates per indicated timepoint in parallel to the gDNA sequencing experiments. RNA isolation, library preparation, and high-throughput sequencing were performed by Singleron Biotechnology (Cologne, Germany) utilizing the plate-based AccuraCode^®^ platform (Singleron Biotechnology). Raw single-end sequencing reads were quality-filtered and adapter-trimmed using fastp, and post-trimming quality was verified via FastQC. High-quality reads were aligned to the mouse reference genome (*Mus musculus*, GRCm39, RRID:SCR_021169) using STAR (version 2.7.10). The average unique mapping rate was 85%, with >95% of reads mapping to exonic regions. rRNA contamination was <2% across all samples. Gene-level expression quantification was executed to generate a raw count matrix containing Ensembl gene IDs, gene symbols, and read counts.

Downstream normalization and differential gene expression analyses were performed in RStudio (version 4.5.1, Posit, RRID:SCR_000432) using the DESeq2 R package (RRID:SCR_015687) via custom scripts. Transcriptomic profiles of the irradiated phases (T1 and T4) were compared against non-irradiated control samples (Control). Statistically significant differentially expressed genes (DEGs) were defined using a threshold of an adjusted *p*-value < 0.05 and an absolute log2 fold-change > 1.

For global expression profiling, clustered heatmaps were generated using the pheatmap R package (RRID:SCR_016418). Genes were pre-filtered to exclude low-expression transcripts with a mean count ≤ 10, and the top 500 highly variable genes were selected based on the highest row variance. Expression values were scaled as row Z-scores. Volcano plots were constructed using the EnhancedVolcano R package (RRID:SCR_018931) combined with ggplot2 and ggrepel.

Functional enrichment was conducted via Gene Set Enrichment Analysis (GSEA) using the clusterProfiler R Package (RRID:SCR_016884), which utilizes the fgsea algorithm (RRID:SCR_020938) for fast enrichment calculations. *Mus musculus* gene sets were retrieved from the Molecular Signatures Database (MSigDB, RRID:SCR_016863) via the msigdbr R package (RRID:SCR_025216). GSEA parameters were restricted to gene sets with a minimum size of 15 and a maximum size of 500 genes. For visualization in bubble plots, pathways were ranked by their significance score, quantified as −log_10_ (adjusted *p*-value), selecting the top 10 upregulated and top 10 downregulated biological processes per comparison.

To ensure reproducibility, all custom scripts utilized for differential expression analysis and functional enrichment are hosted on the Open Science Framework (OSF) repository.

### Invasion assay

A 3D spheroid invasion assay was performed to evaluate the *in vitro* metastatic and invasive potential of parental (non-irradiated) and Rx cells. Cells were seeded at a density of 10,000 cells/well in U-bottom 96-well plates (Greiner, Cat# 650185) to prevent cell attachment and promote formation of a single tridimensional aggregate. The plates were incubated at 37°C with 5% CO_2_ for 24 hours to allow spheroid formation. Following the 24-hour incubation, 70 µl of the culture medium was carefully removed and replaced with 30 µl of Matrigel^®^ (Corning, RRID:SCR_025217) (1:1 final ratio) to create an extracellular matrix environment. Brightfield images were captured 24 hours after Matrigel addition (T_0_) and at day 15 (T_15_). Quantitative image analysis was performed using QuPath software (version 0.7.0, University of Edinburgh, RRID:SCR_018257), and two distinct parameters were calculated for each spheroid: pure growth (determined by the expansion of the core spheroid body), and normalized invasion (quantified as the total invasive area subtracted by the initial T_0_ area and normalized to the baseline growth). All conditions were performed in triplicate.

### *In vivo* experiments

Immunocompetent 129J/B6 F1 mice were utilized to compare the metastatic ability of the parental and the Rx *in vivo*. All animal experiments were approved by the Landesamt für Verbraucherschutz und Ernährung (LAVE, approval numbers 81-02.04.2019.A291 and 2024-509) and conducted in accordance with institutional guidelines. To prevent sex-specific antigen immune rejection, cell lines were injected into mice matching the biological sex of the cell line origin. Sex identity was verified via PCR amplification of *Rbm31x* (269 bp) and *Rbm31y* (353 bp) genes, identifying male-derived lines by two bands and female-derived cell lines by a single band (34). Mice at an age of 9 to 14 weeks were used. Animal weight, behavior, and tumor growth were monitored regularly. Mice were sacrificed upon reaching a cumulative severity score of 20 of when subcutaneous tumors reached a volume of 1000 mm^3^.

#### Subcutaneous and intravenous injections

For subcutaneous assays (n= 5 mice/group), parental and Rx cells (5 x10^4^ cells in 100 µl of sterile PBS) were injected into both flanks in a 1:1 mixture with Matrigel^®^ (Corning). For intravenous injections (n= 3-4 mice/group), cells (1 - 5 x10^4^ cells in 100 µl of sterile PBS) were injected via the tail vein. At the endpoint, tumors and lungs were harvested, weighed, and macroscopically evaluated using a Zeiss SteREO Discovery.V8 microscope (Carl Zeiss Microscopy, Germany, RRID:SCR_021115). For GFP/Tomato-expressing cell lines, two of the five lung lobes were processed for flow cytometry (BD FACSCelesta™, BD Biosciences, RRID:SCR_019597), while the remaining three lobes were inflated with 4% PFA for histology. For non-fluorescent cell lines, all five lung lobes were collected for histological analysis.

#### Lung dissociation and flow cytometry

Harvested lung lobes were minced and enzymatically digested in HBSS (Gibco, Cat# 14175095) supplemented with 0.25% Trypsin-EDTA (Thermo Fisher Scientific, Cat# 25200056), 10 mg/ml Collagenase IV (PanBiotech, Cat# P04-22100), and dispase (Corning, Cat# 354235) for 30 minutes at 37°C under constant rotation. To terminate enzymatic activity, the suspension was cooled on ice for 5 minutes, and a quenching medium composed of Leibovitz L-15 (Thermo Fisher Scientific, Cat# 11415064) supplemented with FBS (VWR/Biowest, Cat# S1810) and 5 mg/ml DNase I (Thermo Fisher Scientific, Cat# EN0521) was added. The cell suspension was filtered through a 40 µm mesh, and red blood cells were lysed using Ammonium-Chloride-Potassium (ACK) buffer. Cells were resuspended in FACS buffer (PBS + 2% FBS) with DAPI (1:1000, Thermo Fisher Scientific, Cat# D1306) for dead-cell exclusion. Data were acquired on a BD FACSCelesta^TM^ flow cytometer (BD Biosciences) using BD FACSDiva^TM^ software (BD Biosciences, RRID:SCR_001456) and analyzed with FlowJoTM software (version 10, FlowJo LLC, RRID:SCR_008520). Representative plots and the flow cytometer gating strategy are shown in **Supplementary Figure S7B**.

#### Histology, immunohistochemistry and quantification

PFA-tissues were dehydrated, embedded in paraffin, and sectioned at 2 µm. Morphological evaluation was performed via Hematoxylin and Eosin (H&E) staining using a Leica Autostainer XL (Leica Biosystems, RRID:SCR_020212). For GFP-expressing cell lines, immunohistochemistry (IHC) was performed using a primary goat anti-GFP antibody (Abcam, 1:500, RRID:AB_305643) and a secondary donkey anti-goat HRP antibody (Abcam, 1:200, RRID:AB_228395). Signal was developed with HRP-DAB (Dako, RRID:AB_2891390) and counterstained with hematoxylin. Slides were imaged at 10x using an Axioscan Z1 slide scanner (Zeiss, RRID:SCR_022858). Metastatic burden was quantified by manually contouring metastases in Fiji (ImageJ v1.53c, NIH, RRID:SCR_003070). The metastatic area was calculated as a percentage of the total tissue area per lobe, and the final load was expressed as the average percentage across all lobes.

### Metabolic assays

To characterize and compare the metabolic profiles of parental and Rx cell lines, intracellular oxidative stress, mitochondrial respiration, and total ATP production were evaluated.

For flow cytometry assays, specific fluorogenic probes were used, and all incubation steps performed at 37°C. Total intracellular ROS production was assessed by incubating the cells with either 500 nM CellROX™ Orange (Invitrogen, Cat# C10493 ) or Deep Red (Invitrogen, Cat# C10491) for 60 min, or with 1 μM CM-H2DCFDA (Thermo Scientific, Cat# C6827) for 30 min, depending on the fluorophore expression profile of each cell line (GFP, GFP and Tomato, or Tomato-only, respectively). Hydrogen peroxide (H₂O₂, 1000 μM, Carl Roth, Cat# 8070) served as a positive control in all ROS assays. Representative plots and the flow cytometer gating strategy are shown in **Supplementary Figure S7C**. Mitochondrial ROS levels were determined by incubating cells for 30 min with either 5 μM MitoSOX™ Red (MedChemExpress, Cat# 1003197-00-9; for GFP-expressing cell lines) or 1 μM MitoSOX™ Green (Invitrogen, Cat# M36006; for Tomato-expressing cell lines) diluted in warm HBSS (Gibco). Representative plots and the flow cytometer gating strategy are shown in **Supplementary Figure S7D**. Mitochondrial membrane potential (MMP) was evaluated using 10 nM Tetramethylrhodamine Methyl Ester (TMRM; Invitrogen) for 30 min; where indicated, cells were pre-treated with 20 μM carbonyl cyanide 4- (trifluoromethoxy) phenylhydrazone (FCCP, MedChem Express, Cat# 370-86-5) for 5 min to validate probe responsiveness. Representative plots and the flow cytometer gating strategy are shown in **Supplementary Figure S7E**. Fluorescence intensities across all assays were acquired on a BD FACSCelesta™ flow cytometer (BD Biosciences) using BD FACSDiva™ software (BD Biosciences, RRID:SCR_019597), and data analysis was performed using FlowJo™ software (FlowJo LLC, RRID:SCR_008520).

Mitochondrial respiration was quantified using a Seahorse XFe96 Extracellular Flux Analyzer (Agilent Technologies, RRID:SCR_019545) via the XF Cell Mito Stress Test (Agilent Technologies, RRID:SCR_026230). Oxygen consumption rate (OCR) was monitored baseline and following sequential injections of Oligomycin (1.5 μM), FCCP (1 μM), and Rotenone/Antimycin A (0.5 μM). All data were normalized to cell confluency via NYONE (Synentec GmbH) and analyzed using the Seahorse Wave Software (Agilent Technologies, RRID: SCR_016952).

Total ATP production levels were measured using the CellTiter-Glo^®^ 2.0 Assay (Promega, RRID:SCR_025211). Cells were seeded in 96 well F-bottom (Chimney well) µClear white plate (Greiner) at a density of 10,000 cells/well and incubated for 7 hours. Following the manufacturer’s instructions with slight modifications, the CellTiter-Glo^®^ 2.0 reagent was diluted 1:2 in PBS and added to the wells in a 1:1 ratio relative to the culture medium. Luminescence was subsequently measured using a TECAN Spark multimode reader (TECAN Trading AG, Switzerland, RRID:SCR_024614) with an integration time of 1 second.

### Determination of mitochondrial DNA copy number by real-time PCR

To evaluate the relative mitochondrial DNA (mtDNA) content per cell, the ratio between mtDNA and nuclear DNA (nDNA) was determined via quantitative real-time PCR (qPCR) in parental and Rx cell lines. Reactions were performed using 5 ng/µl of gDNA as a template to amplify the mitochondrial *Nd4* gene and the nuclear *K19* gene, respectively. Amplifications were executed in LightCycler^®^ 480 Multiwell Plate 96 (Roche, Cat# 04729692001) plates using a LightCycler^®^ 480 II instrument (Roche, RRID:SCR_025134). Relative mtDNA copy number was calculated using the 2^−ΔΔCt^ method by normalizing *Nd4* levels to the *K19* nuclear reference control.

### Statistical analysis

All *in vitro* experiments were performed in three independent biological replicates (N=3) with each three technical replicates (n=3). Results were expressed as mean ±SEM. Statistical differences between parental and Rx cell lines were determined using an unpaired Student’s t-test. For colony formation assays, an ordinary two-way ANOVA followed by Šídák’s multiple comparisons test was performed. The statistical significance between groups for the *in vivo* experiments was determined using a Mann-Whitney test. All analysis and graphing were performed using GraphPad Prism (version 9.5.1, GraphPad Software, RRID:SCR_002798).

## Supporting information

Martinez-Cruz et al. Supplementary Figures S1-S7

## KEY RESOURCES TABLE

| **REAGENT or RESOURCE** | **SOURCE** | **IDENTIFIER** |
|---|---|---|
| **Antibodies** |  |  |
| Anti-GFP antibody | Abcam | Cat# ab6673, RRID:AB_305643 |
| Rabbit Anti-Goat H&L (HRP) | Invitrogen | Cat# 31402, RRID:AB_228395 |
| **Enzymes and Matrices** |  |  |
| Collagenase Type IV | Thermo Fisher Scientific | Cat# 17104019 |
| Dispase II | Corning | Cat# 354235 |
| Dnase I | Thermo Fisher Scientific | Cat# EN0521 |
| Matrigel<sup>®</sup> | Corning | Cat# 356231, RRID:SCR_025217 |
| **Fluorescent probes and dyes** |  |  |
| CellROX Orange<sup>TM</sup> | Invitrogen | Cat# C10493 |
| CellROX Deep Red<sup>TM</sup> | Invitrogen | Cat# C10491 |
| CM-H2DCFDA | Invitrogen | Cat# C6827 |
| mitoSOX<sup>TM</sup> Red | MedChem Express | Cat# HY-D1055 |
| mitoSOX<sup>TM</sup> Green | Invitrogen | Cat# M36006 |
| Tetramethylrhodamine, Methyl Ester, Perchlorate (TMRM) | Thermo Fisher Scientific | Cat# T668 |
| DAPI | Thermo Fisher Scientific | Cat# D1306 |
| GelRed<sup>®</sup> | Sigma-Aldrich | Cat# S9812 |
| **Chemicals and experimental controls** |  |  |
| Carbonyl cyanide 4-(trifluoromethoxy)phenylhydrazone (FCCP) | MedChem Express | Cat# HY-100410 |
| Antimycin A | Merck | Cat# A8674 |
| Hydrogen peroxide (H<sub>2</sub>O<sub>2</sub>) 30% | Carl Roth | Cat# 8070 |
| **Molecular biology reagents** |  |  |
| KAPA HiFi HotStart ReadyMix | Roche | Cat# 07958935001 |
| SYBR<sup>TM</sup> Green Master Mix | Applied Biosystems | Cat# 4309155 |
| **Equipment** |  |  |
| Cytation 5 Imaging Multi-Mode Reader | Agilent Technologies | RRID:SCR_019732 |
| NYONE automated cell imager | Synentec GmbH | RRID: SCR_024623 |
| RS320 Cabinet Irradiator | Xstrahl, Ltd. | RRID:SCR_024344 |
| **Deposited data** |  |  |
| DNA-seq barcoded cell lines (394T4, 482N1, 579T1, 579DLN) | This paper | Bioproject number: PRJNA1492111 |
| RNA-seq barcoded (394T4, 482N1, 579T1, 579DLN) and non-barcoded (368T1, 238N1) cell lines | This paper | Bioproject number: PRJNA1492111<br>GEO record number: GSE337825 |
| **Experimental models: Cell lines** |  |  |
| Murine 394T4 | Winslow et al. (9) | N/A |
| Murine 368T1 | Winslow et al. (9) | N/A |
| Murine 579T1 | Winslow et al. (9) | N/A |
| Murine 482N1 | Winslow et al. (9) | N/A |
| Murine 238N1 | Winslow et al. (9) | N/A |
| Murine 579DLN | Winslow et al. (9) | N/A |
| HEK293T packaging cells | ATCC | #CRL-11268 |
| **Experimental models: Organisms/strains** |  |  |
| Immunocompetent 129J/B6 F1 mice | The Jackson Laboratory via Charles River | Cross of #002448 (RRID:IMSR_JAX:002448) and #000664 (RRID:IMSR_JAX:000664) |
| **Oligonucleotides** |  |  |
| DNA-sequencing primers, qPCR primers | Eurofins Genomics | See Supplementary Table 1 |
| **Recombinant DNA** |  |  |
| pMD2.G | Addgene | #12259 |
| pMDLg/pRRE | Addgene | #12251 |
| pRSV-Rev | Addgene | #12253 |
| TransIT®-Lenti Transfection Reagent | Mirus Bio | #MIR 6600 |
| Lentiviral transfer vector pLenti CMV GFP Hygro 9N | This paper | N/A |
| **Software** |  |  |
| Affinity Designer 2 | Serif Europe | RRID:SCR_016952 |
| Agilent Seahorse Analytics/Wave | Agilent Technologies | RRID:SCR_024491 |
| Gen 5<sup>TM</sup> Microplate Reader and Imager Software | Agilent Technologies | Version 3.14, RRID:SCR_017317 |
| Graphpad Prism | Graphpad Software | Version 9.5.1, RRID:SCR_002798 |
| RStudio | Posit | Version 4.5.1, RRID:SCR_000432 |
| DESeq2 | Bioconductor | RRID:SCR_015687 |
| EnhancedVolcano | Bioconductor | RRID:SCR_018931 |
| pheatmap | Bioconductor | RRID:SCR_016418 |
| clusterProfiler | Bioconductor | RRID:SCR_016884 |
| fgsea | Bioconductor | RRID:SCR_020938 |
| msigdbr | Bioconductor | RRID:SCR_025216 |
| QuPath | University of Edinburgh | v0.7.0, RRID:SCR_018257 |
| SnapGene | Dotmatics | RRID:SCR_015053 |
| Synentec cell recognition software | Synentec GmbH | RRID: SCR_024623 |
| FlowJo V10 | FlowJo LLC | RRID:SCR_008520 |
| ImageJ/Fiji | NIH | RRID:SCR_003070 |
| Python | Python Software Foundation | Version 3.9.9<br>RRID:SCR_008394 |
| Zen | Zeiss | Version 3.5, RRID:SCR_013672 |

---

**Supplementary Table 1.**
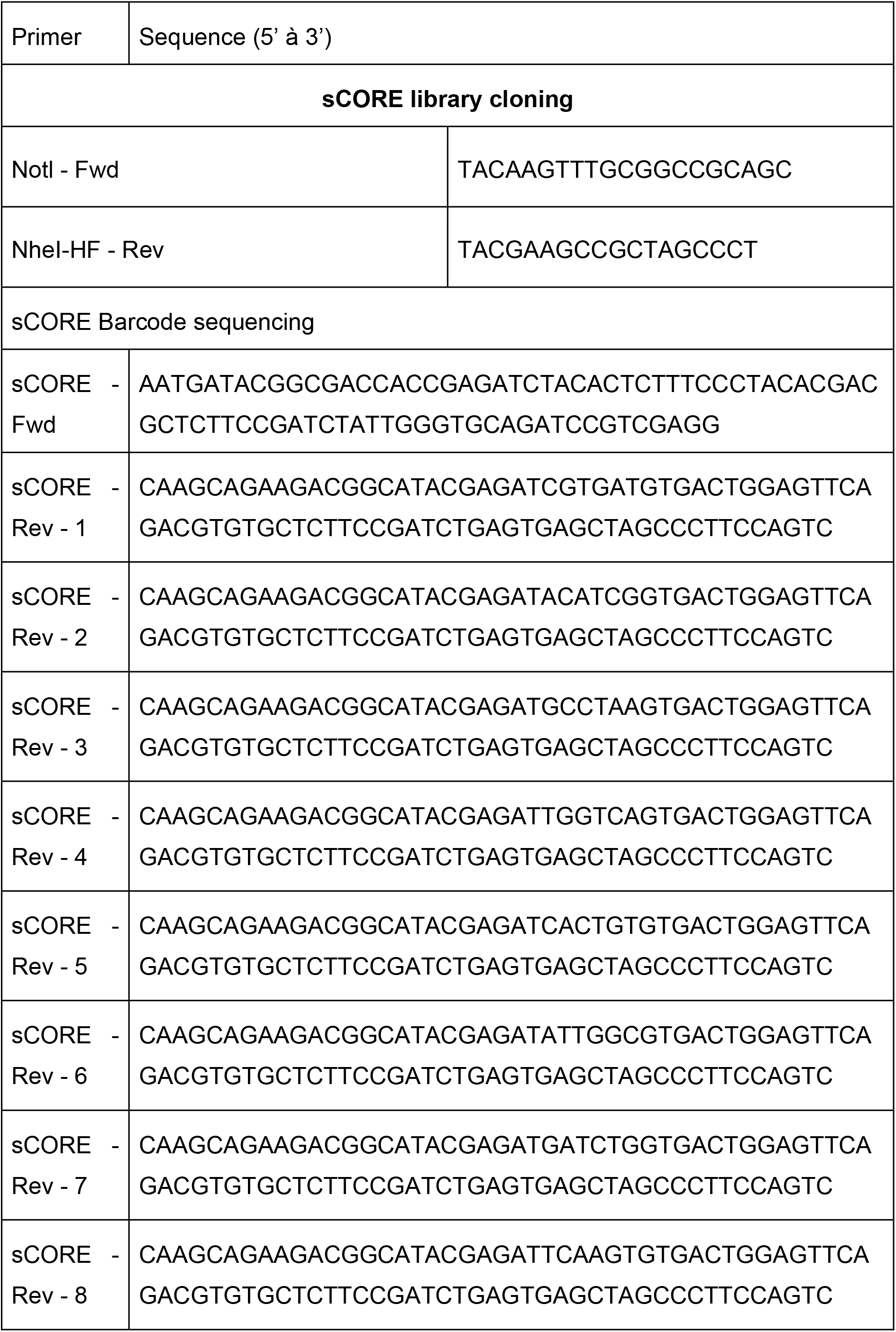
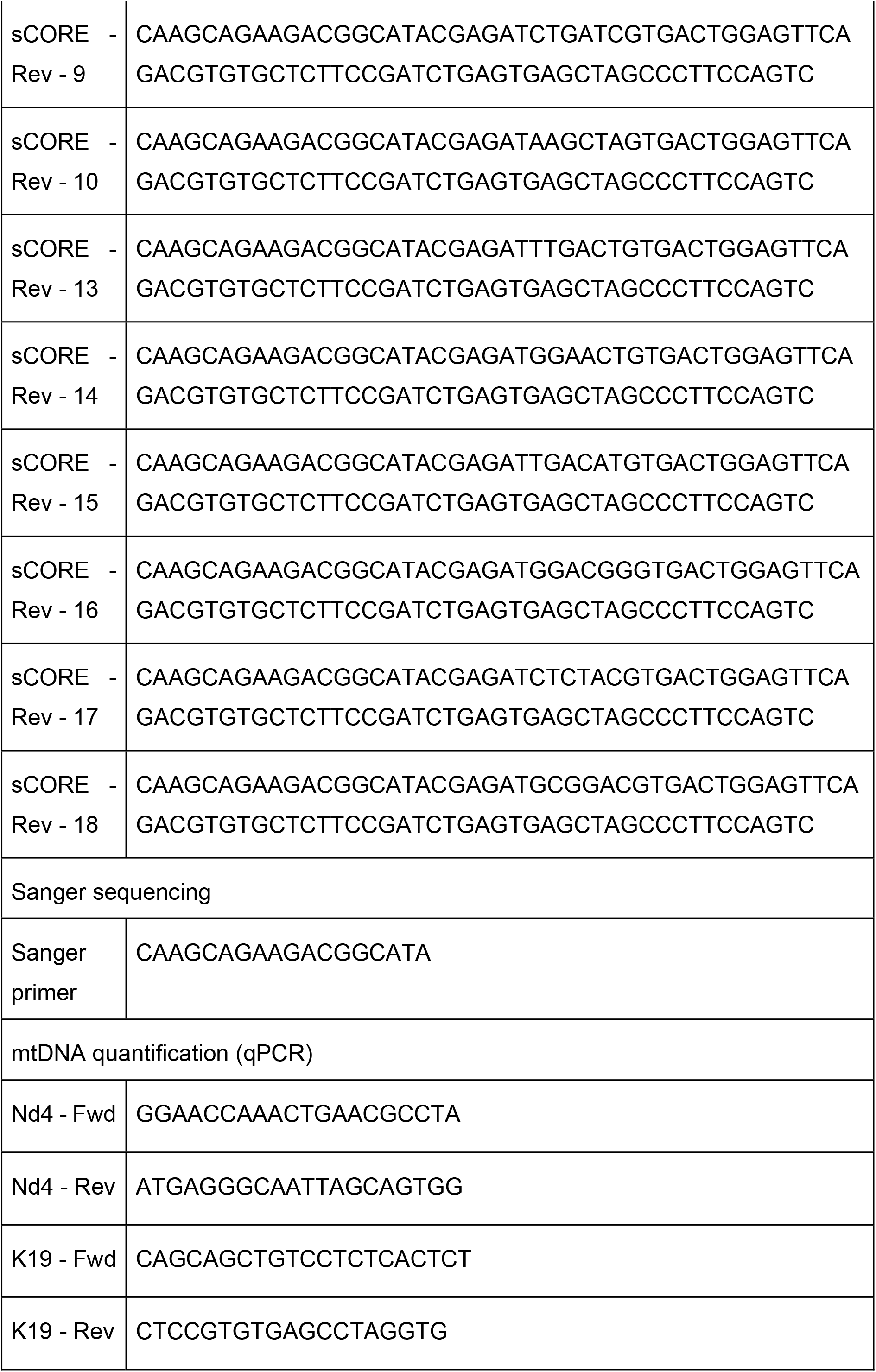
List of oligonucleotide primer sequences used for sCORE barcoding sequencing, Sanger sequencing, and qPCR-based mtDNA quantification.

## ACKNOWLEDGEMENTS

We thank Eva Hahn and Nancy Meyer for excellent technical assistance. We thank the members of the Grüner, Grünwald, and Schramm labs for helpful comments and discussions. We also thank the central animal facility of the University Hospital Essen (ZTL) for their support. This work was funded by the Deutsche Forschungsgemeinschaft (DFG, German Research Foundation) – GRK2762 – project number 450917483 (projects L1 to B.M.G., L2 to J.M., L3 to A.S., T1 to V.J., and M1 to D.H.).

## AUTHOR CONTRIBUTIONS

Conceptualization: B.M.G.

Methodology: M.M.C., M.D., A.B, D.H., B.M.G.

Formal analysis: M.M.C., M.D., D.H., S.L.

Investigation: M.M.C., V.D., V.A., J.J

Visualization: M.M.C., M.D.

Resources, concepts, and funding acquisition: J.M., V.J., A.S., D.H., B.M.G.

Writing – original draft: M.M.C., B.M.G.

Writing – review & editing: all authors

