## Supplementary material for "Radiation survival selects clonally restricted NSCLC survivors with impaired metastatic capacity and distinct bioenergetic states": Martinez-Cruz et al. Supplementary Figures S1-S7

**Supplementary Figure 1. Heterogeneous radiosensitivity and clonal tracking setup**

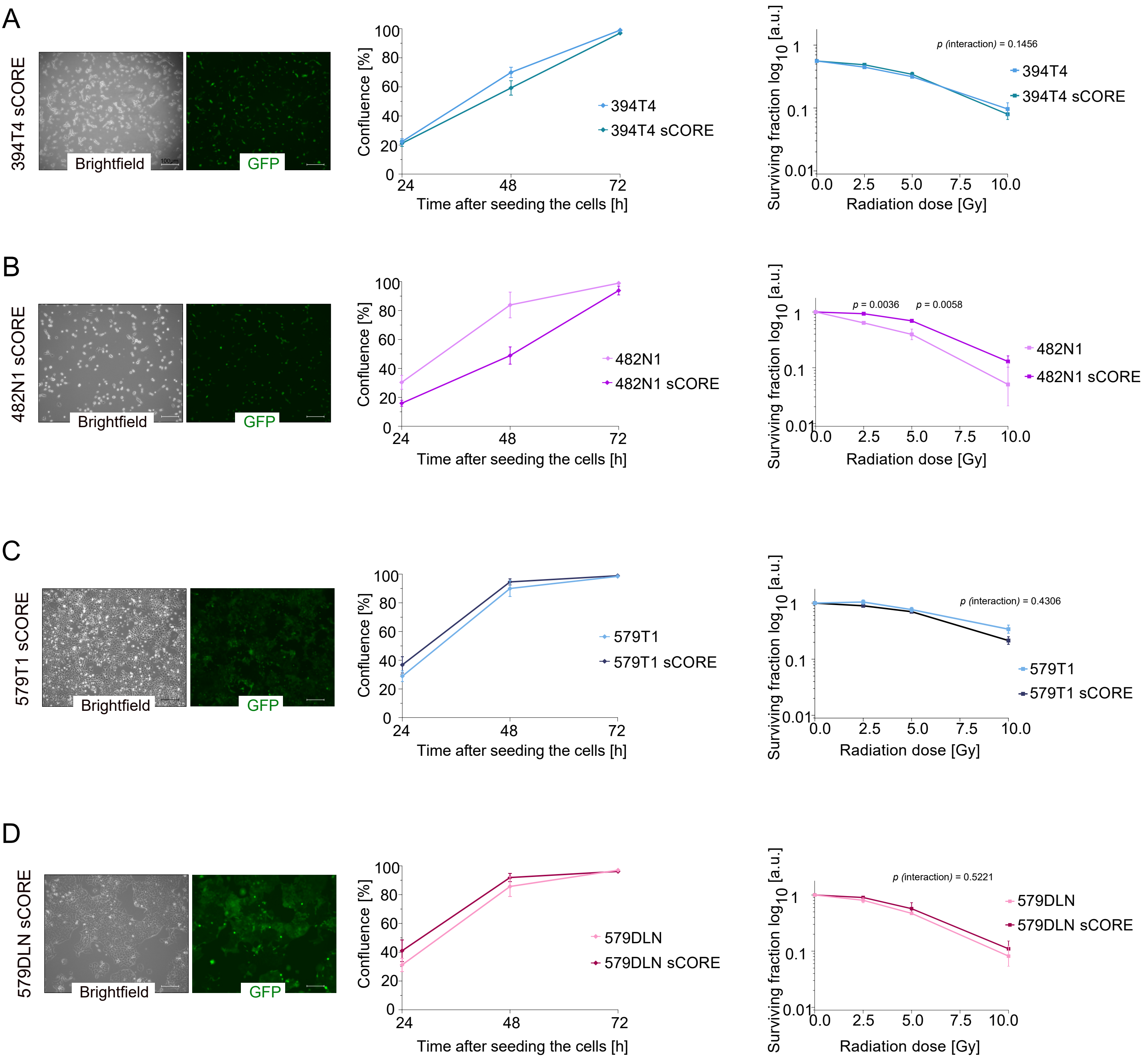

**Supplementary Figure S1. Heterogeneous radiosensitivity and clonal tracking setup. (A–D)** Evaluation of the sCORE system in (**A**) 394T4, (**B**) 482N1, (**C**) 579T1, and (**D**) 579DLN. Left panels: representative brightfield (BF) and fluorescence microscopy images showing successful barcoding via GFP expression after lentiviral transduction, sorting, and enrichment. Scale bar: 100 μm. Middle panels: proliferation kinetics of barcoded (sCORE) vs. parental cells over 72 hours. Data are mean ± SEM from three independent biological replicates (N=3). Right panels: clonogenic survival curves after irradiation (0–10 Gy). Data are mean ± SEM from N=3 independent experiments.

**Supplementary Figure 2, page 1/2. Radiation survival is a clonal event that reveals two distinct clonal trajectories**

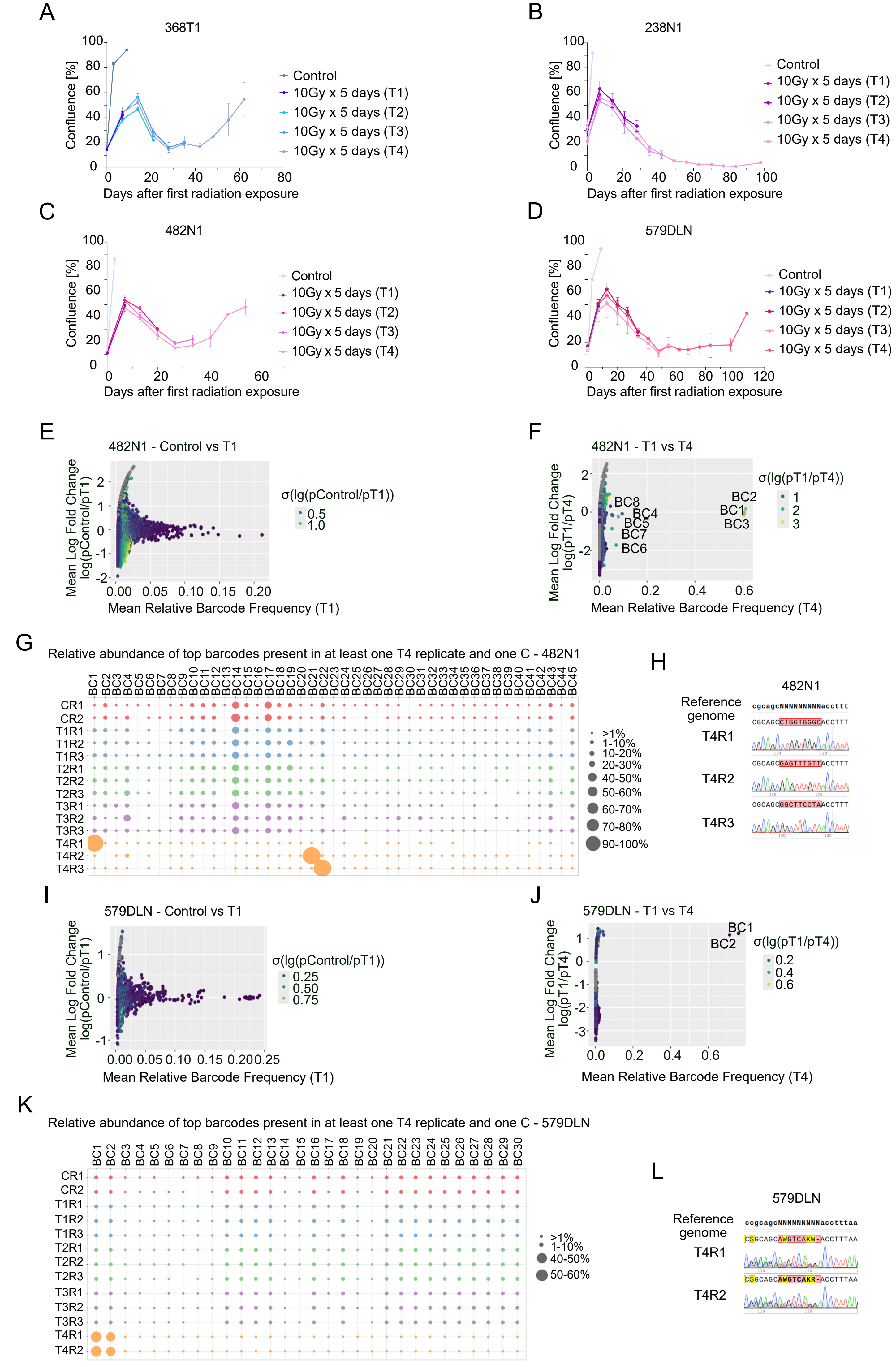

**Supplementary Figure 2, page 2/2. Radiation survival is a clonal event that reveals two distinct clonal trajectories**

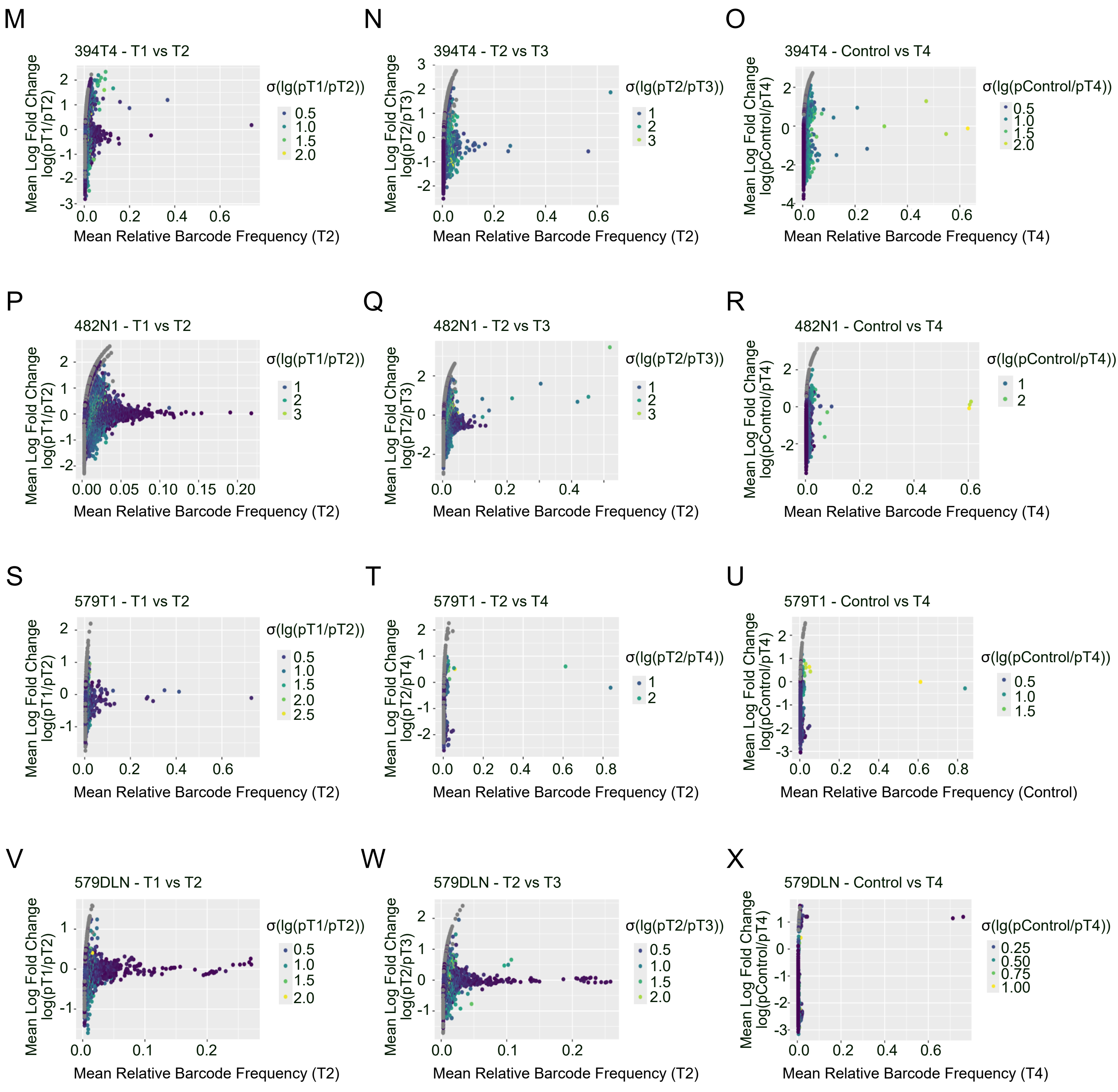

**Supplementary Figure S2. Radiation survival is a clonal event that reveals two distinct clonal trajectories. (A-D)** Proliferation curves, measured as percentage of confluence, for the parental, unbarcoded cell lines **(A)** 368T1, **(B)** 238N1, and the barcoded cell lines **(C)** 482N1 and **(D)** 579DLN under radiation as indicated in main Figure 2A. Color coding distinguishes the non-irradiated control from the populations harvested at different time points (T1-T4). Data points represent mean values ± SEM. **(E, F, I, J, M-X)** Scatter plots of relative barcode frequencies between time points for (**M–O**) 394T4, (**E–F, P–R**) 482N1, (**S–U**) 579T1, and (**I–J, V–X**) 579DLN. Panels show: (**E, I)** Control vs. T1, (**F, J**) T1 vs. T4, (**M, P, S, V**) T1 vs. T2, (**N, Q, W**) T2 vs. T3, (**T**) T2 vs. T4. $Log_2$ fold change is plotted against mean barcode abundance. **(G, K)** Bubble plots depicting the relative abundance of the top barcodes present in at least one T4 replicate and one control replicate for **(G)** 482N1 and **(K)** 579DLN. Color coding distinguishes the different time points, while bubble size corresponds to relative abundance (%). **(H, L)** Sanger chromatograms of n=2 or 3 technical replicates aligned to the reference genome for **(H)** 482N1 and **(L)** 579DLN. Within the replicates, the barcode region is highlighted in red to denote a complete match, whereas mismatches are indicated in yellow. In the reference genome, the barcode region is represented as NNNNNNNNN.

**Supplementary Figure 3, page 1/1. Radiation survivors exhibit increased radiosensitivity and impaired metastatic ability**

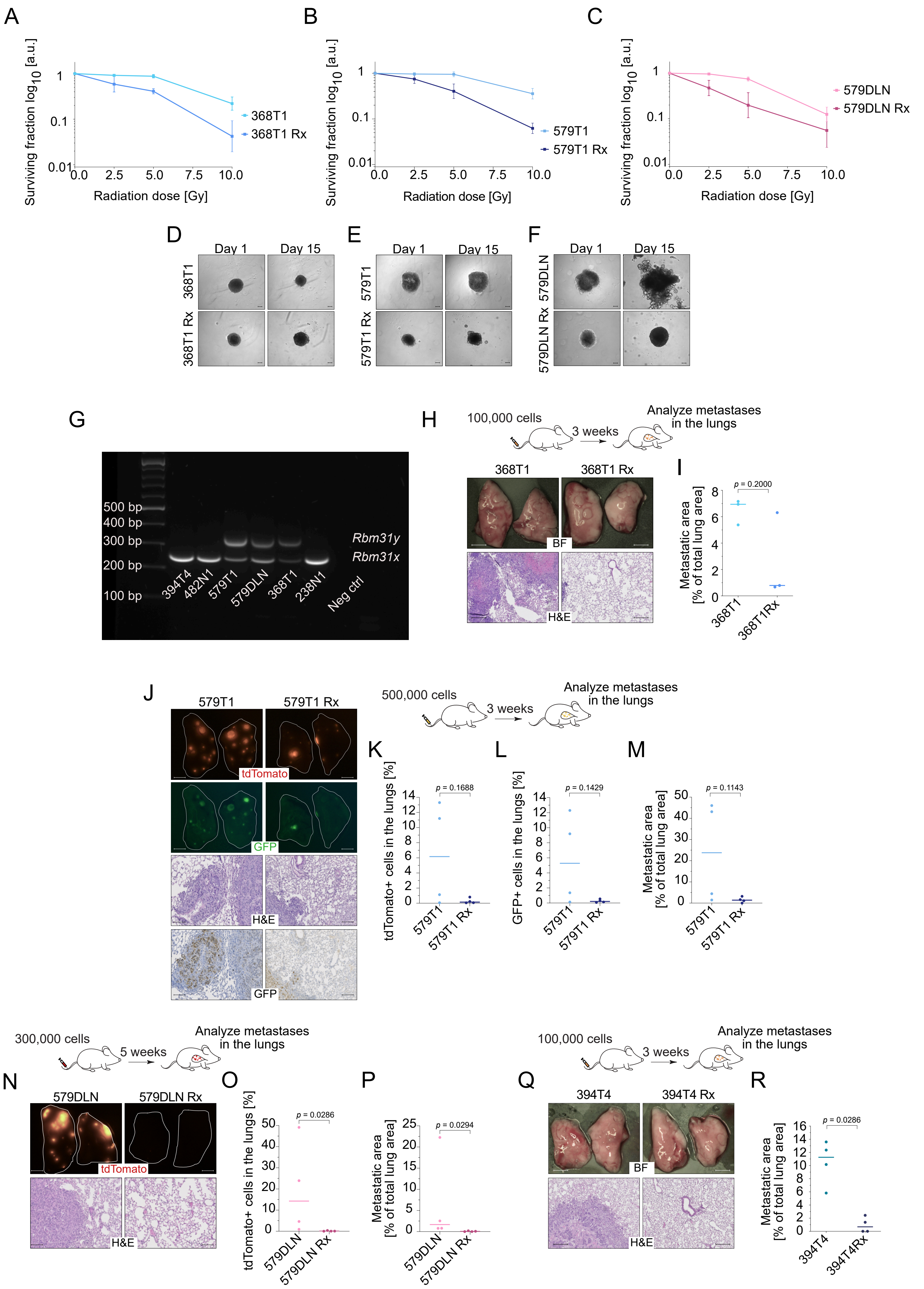

**Supplementary Figure S3. Radiation survivors exhibit increased radiosensitivity and impaired metastatic ability. (A–C)** Radiation-induced clonogenic survival curves of parental versus radiation-surviving (Rx) counterparts for **(A)** 368T1, **(B)** 579T1, and **(C)** 579DLN cell lines. Data represent Mean ± SEM from N=3 independent biological replicates. **(D–F)** Spheroid invasion assay in Matrigel® displaying representative brightfield images at days 1 and 15 post-embedding for **(D)** 368T1, **(E)** 579T1, and **(F)** 579DLN pairs. Scale bar: 200 µm. **(G)** PCR gel electrophoresis for *Rbm31x/y* transcripts to determine the sex origin of the different NSCLC cell lines. **(H, I)** Intravenous tail-vein injection model for parental and radiation-surviving (Rx) 368T1 cells. **(H)** Representative images of lungs (top) and H&E-stained lung sections (bottom). **(I)** Histological scoring of total metastatic burden. **(J–M)** Intravenous tail-vein injection model for the 579T1 pair. **(J)** Representative fluorescence images of lungs showing tdTomato$^+$ and GFP$^+$ macroscopic metastases (top panels), with corresponding H&E (middle panels) and anti-GFP IHC (bottom panels) histological sections (Scale bars: 100 µm). **(K, L)** Quantification of disseminated **(K)** tdTomato$^+$ and **(L)** GFP$^+$ tumor cells in lungs by flow cytometry. **(M)** Histological quantification of metastatic area relative to total lung area. **(N–P)** Intravenous tail-vein injection model for parental and radiation-surviving (Rx) 579DLN cells. **(N)** Representative fluorescence images of lungs showing tdTomato$^+$ (top) and H&E-stained lung sections (bottom) (Scale bars: 100 µm). **(O)** Quantification of tdTomato$^+$ cells in lungs by flow cytometry. **(P)** Histological scoring of total metastatic burden. **(Q, R)** Intravenous tail-vein injection model for the non-barcoded 394T4 pair. **(Q)** Representative images of lungs (top) and H&E-stained lung sections (bottom) (Scale bars: 100 µm). **(R)** Histological scoring of total metastatic burden. All dot plots display individual biological replicates (N=3 or 4), horizontal lines represent Mean ± SEM. *p*-values were determined by two-tailed Mann-Whitney U test.

**Supplementary Figure 4 , page 1/5. Radiation survivors converge on proliferative recovery through divergent transcriptional trajectories**

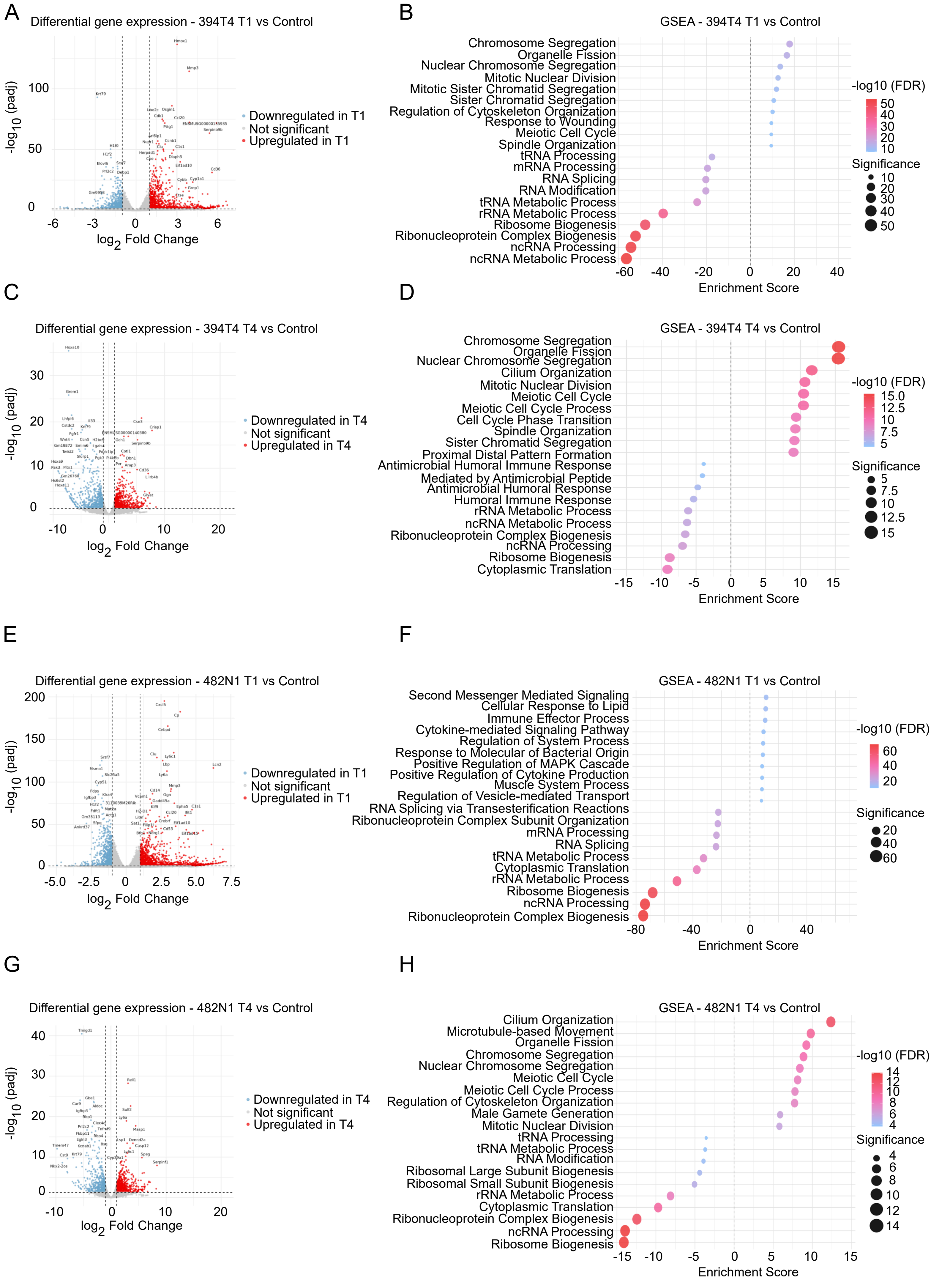

**Supplementary Figure 4, page 2/5. Radiation survivors converge on proliferative recovery through divergent transcriptional trajectories**

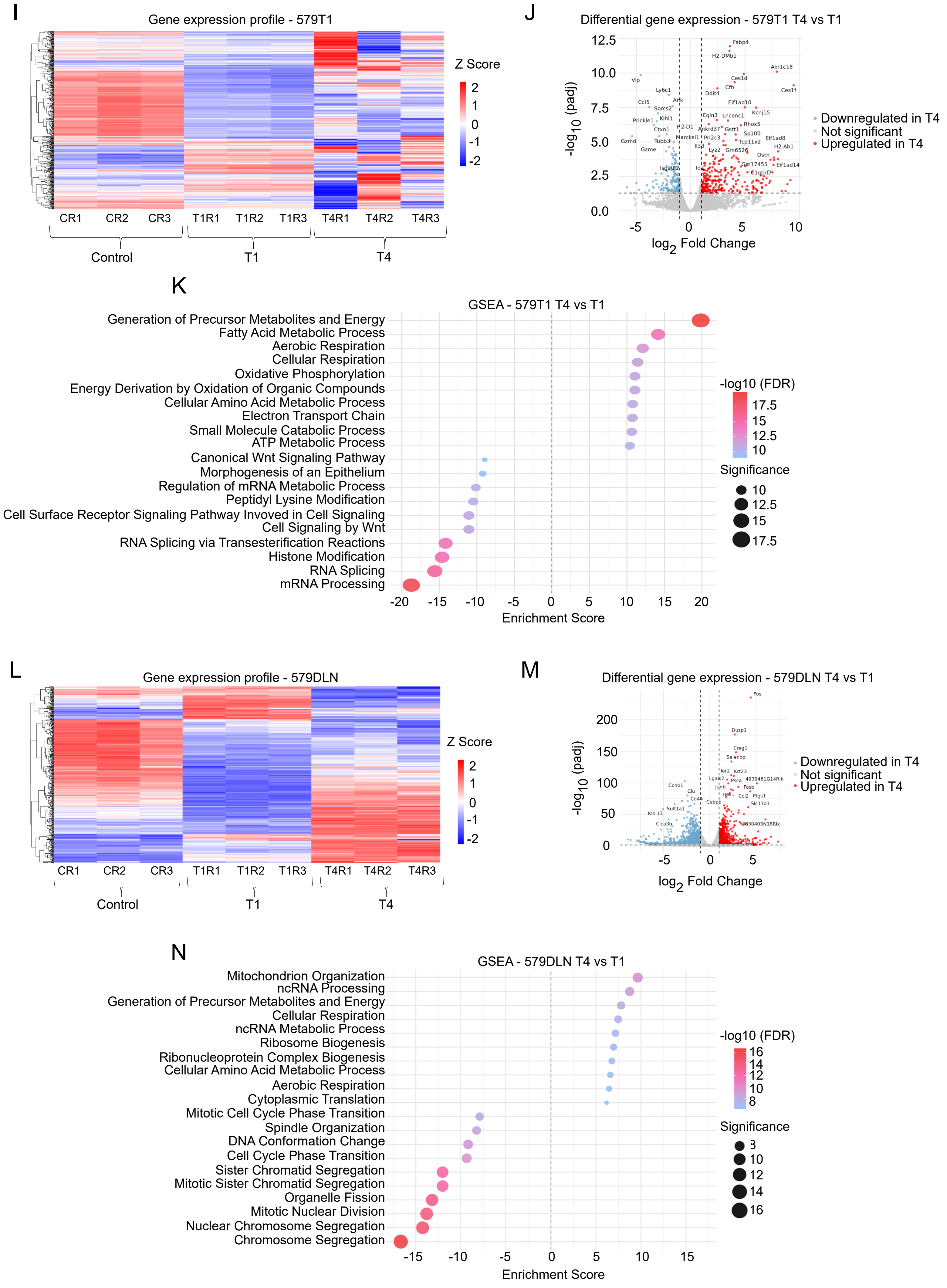

**Supplementary Figure 4, page 3/5. Radiation survivors converge on proliferative recovery through divergent transcriptional trajectories**

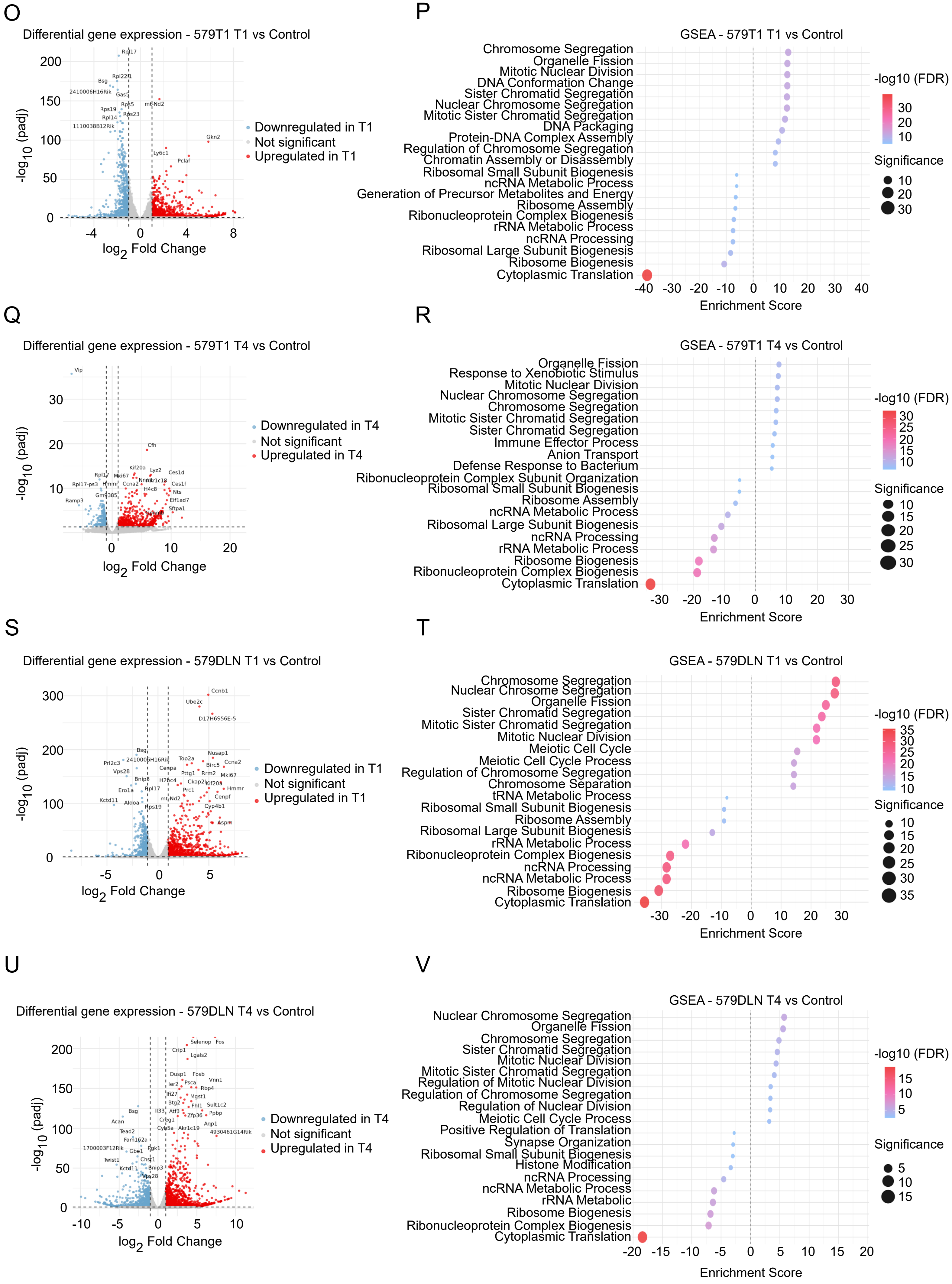

**Supplementary Figure 4, page 4/5. Radiation survivors converge on proliferative recovery through divergent transcriptional trajectories**

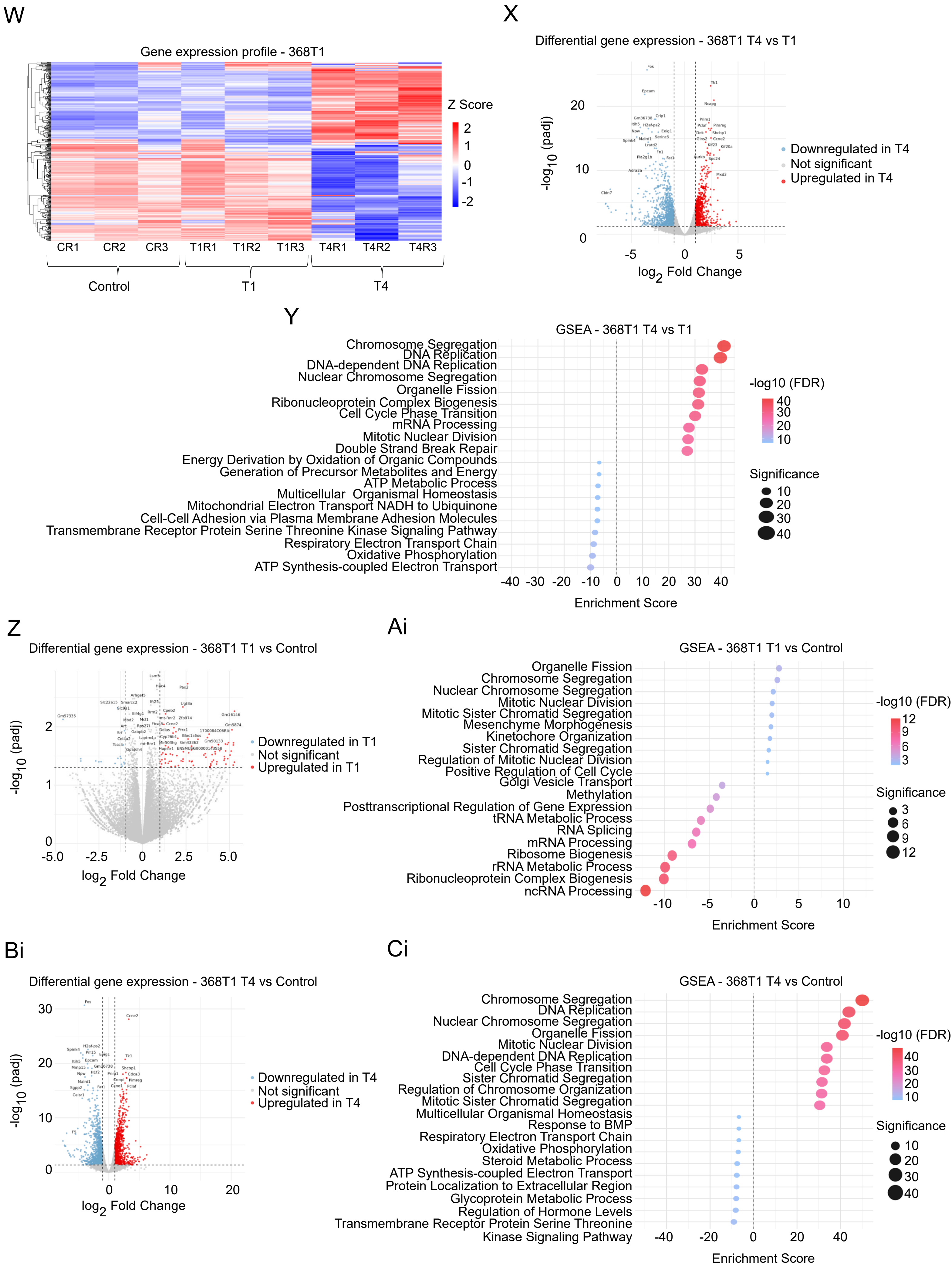

**Supplementary Figure 4, page 5/5. Radiation survivors converge on proliferative recovery through divergent transcriptional trajectories**

Di

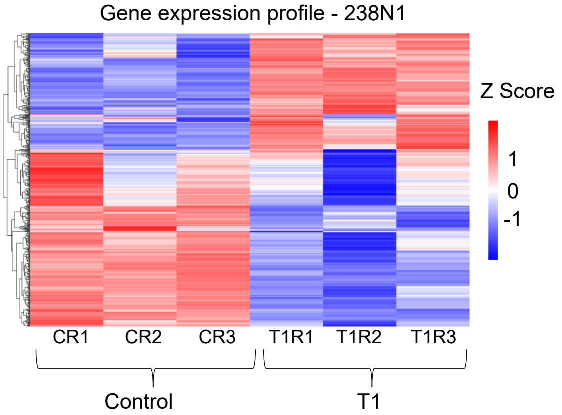


Ei

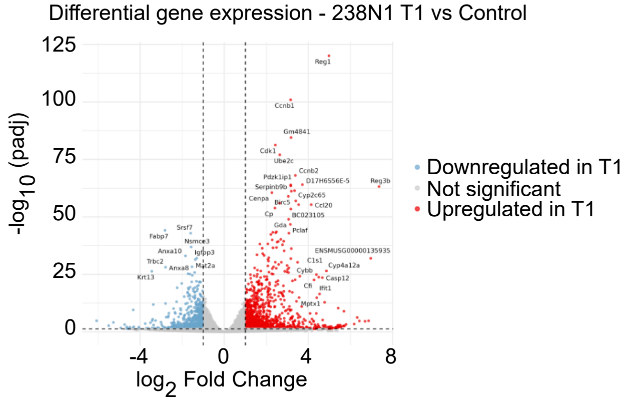


Fi

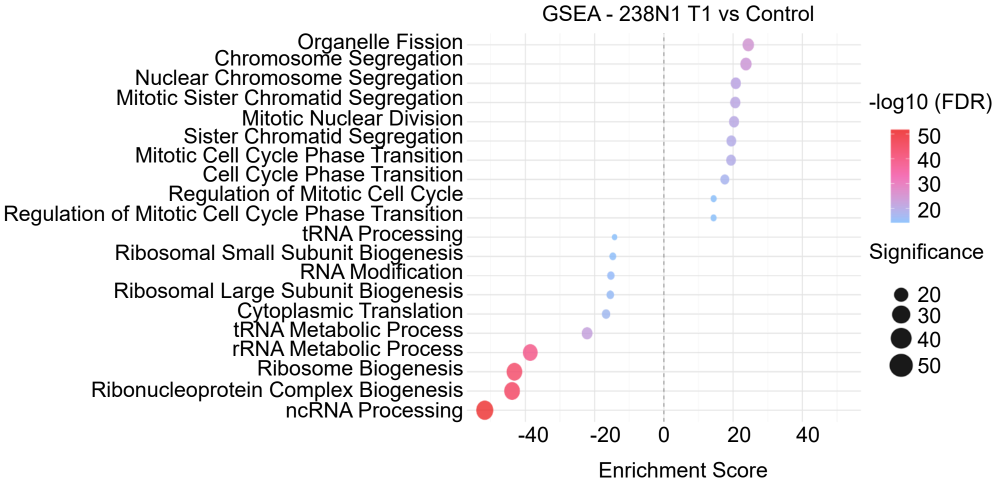

**Supplementary Figure 4. Radiation survivors converge on proliferative recovery through divergent transcriptional trajectories. (A, E, O, S, Z, Ei)** Volcano plots illustrating differentially expressed genes (DEGs) in the acute phase (T1) relative to the control for **(A)** 394T4, **(E)** 482N1, **(O)** 579T1, **(S)** 579DLN, **(Z)** 368T1, and **(Ei)** 238N1 cell lines. **(C, G, Q, U, Bi)** Volcano plots illustrating differentially expressed genes (DEGs) in the recovery phase (T4) relative to the control for **(C)** 394T4, **(G)** 482N1, **(Q)** 579T1, **(U)** 579DLN, and **(Bi)** 368T1 cell lines. **(J, M, X)** Volcano plots illustrating differentially expressed genes (DEGs) in the recovery phase (T4) relative to the acute phase (T1) for **(J)** 579T1, **(M)** 579DLN, and **(X)** 368T1 cell lines. For all volcano plots, dashed vertical and horizontal lines indicate the $\log_2$ fold-change > 1 and statistical significance thresholds ($p_{adj} < 0.05$), respectively. Statistically significant upregulated genes are highlighted in red, whereas downregulated genes are shown in blue. Note that for panel **(Z)**, nominal *p*-values are displayed on the y-axis, whereas adjusted *p*-values ($p_{adj}$) are utilized for all other cell lines. **(B, F, P, T, Ai, Fi)** Gene Set Enrichment Analysis (GSEA) bubble plots displaying significantly enriched biological processes in the acute phase (T1) relative to the control for **(B)** 394T4, **(F)** 482N1, **(P)** 579T1, **(T)** 579DLN, **(Ai)** 368T1, and **(Fi)** 238N1 cell lines. **(D, H, R, V, Ci)** Gene Set Enrichment Analysis (GSEA) bubble plots displaying significantly enriched biological processes in the recovery phase (T4) relative to the control for **(D)** 394T4, **(H)** 482N1, **(R)** 579T1, **(V)** 579DLN, and **(Ci)** 368T1 cell lines. **(K, N, Y)** Gene Set Enrichment Analysis (GSEA) bubble plots displaying significantly enriched biological processes in the recovery phase (T4) relative to the acute phase (T1) for **(K)** 579T1, **(N)** 579DLN, and **(Y)** 368T1 cell lines. For all GSEA plots, the x-axis indicates the normalized enrichment score (NES) for each gene set, where positive and negative values represent enrichment and depletion, respectively. Both bubble size and color intensity reflect the statistical significance scale, quantified as $-\log_{10}$ (FDR), shifting from low significance (blue) to high significance (red). Hierarchical unsupervised clustering heatmaps displaying the top 500 highly variable genes for **(I)** 579T1, **(L)** 579DLN, **(W)** 368T1, and **(Di)** 238N1 cell lines. Genes were pre-filtered to exclude low expression transcripts (mean expression < 10) and selected based on the highest row variance. Subpopulations were analyzed at baseline (Control; CR1–CR3), and at acute (T1; T1R1–T1R3) and recovery phases (T4; T4R1–T4R3), where "R" denotes independent biological replicates (note: 238N1 was only analyzed at Control and T1 time points). Color intensity represents relative gene expression levels scaled as Z-scores, shifting from downregulated (blue) to upregulated (red) states.

**Supplementary Figure 5. Radiation survival is associated with distinct bioenergetic states. (A-C)** Oxygen consumption rate (OCR) kinetics, normalized to cell confluence, measured over time under baseline conditions and following successive injections of Oligomycin, FCCP, and Rotenone/Antimycin A (Rot/AA) for **(A)** 368T1, **(B)** 579T1, and **(C)** 579DLN cell lines. **(D-F)** Basal respiration rates, **(G-I)** spare respiratory capacity, and **(J-L)** coupling efficiency (%) for **(D, G, J)** 368T1, **(E, H, K)** 579T1, and **(F, I, L)** 579DLN. Seahorse kinetic data represent n=4 technical replicates within one out of three biological replicates (N=3), bar graphs represent Mean ± SEM from N=3 independent biological replicates. *p*-values were determined using an unpaired Student's t-test with Welch's correction.

**Supplementary Figure 6, page 1/1. Radiation survival is associated with distinct bioenergetic states**

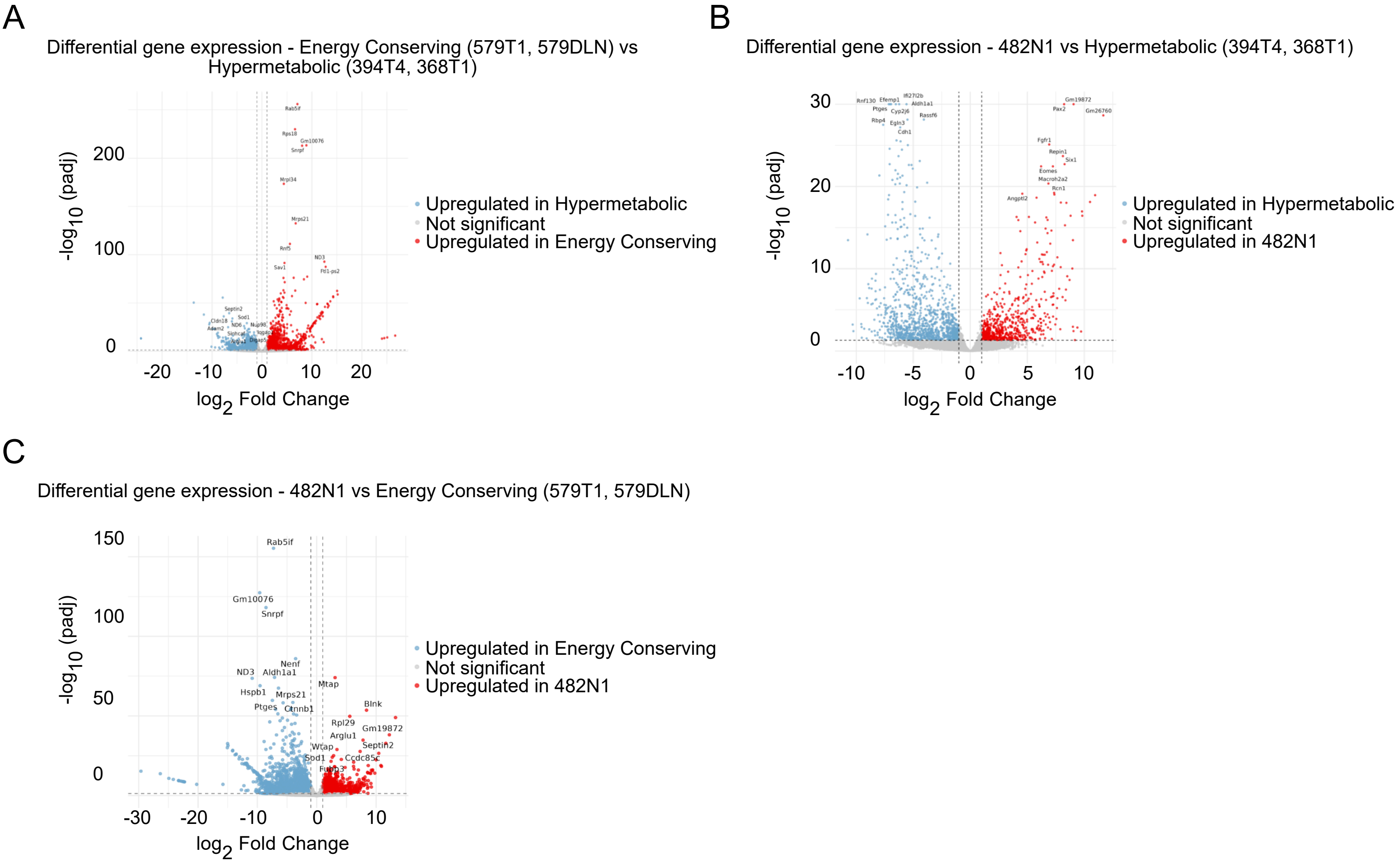

**Supplementary Figure 6. Radiation survival is associated with distinct bioenergetic states. (A-C)** Volcano plots illustrating differentially expressed genes (DEGs) for distinct phenotypic comparisons: **(A)** Energy-conserving cell lines (579T1, 579DLN) compared against Hypermetabolic cell lines (394T4, 368T1), **(B)** 482N1 compared against Hypermetabolic cell lines (394T4, 368T1), and **(C)** 482N1 compared against Energy-conserving cell lines (579T1, 579DLN). For all volcano plots, dashed vertical and horizontal lines indicate the $\log_2$ fold-change >1 and statistical significance thresholds (*p*-adjusted < 0.05), respectively. Statistically significant upregulated genes are highlighted in red, whereas downregulated genes are shown in blue.

**Supplementary Figure 7, page 1/1**

A

Barcode transduction and purity

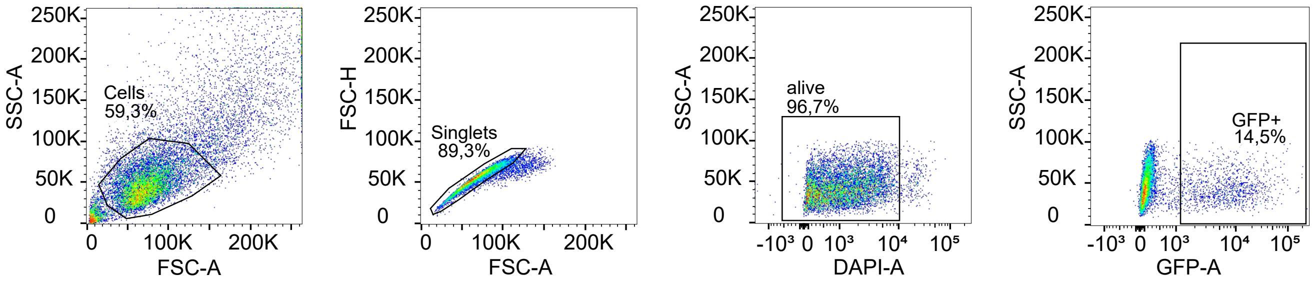


B

Lung metastasis quantification

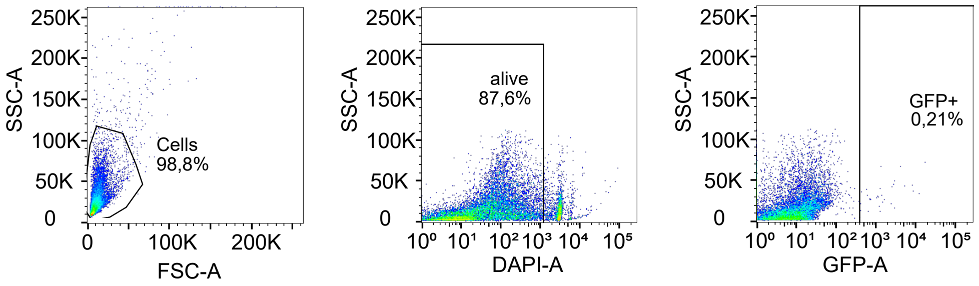


C

Cytosolic ROS

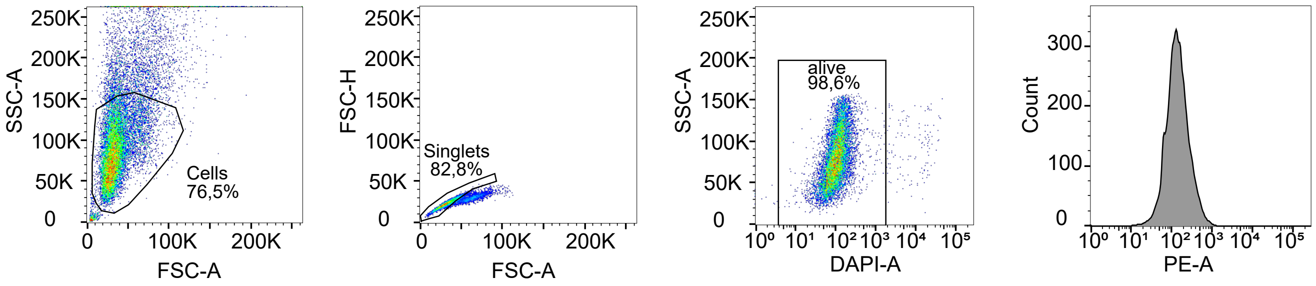


D

Mitochondrial ROS

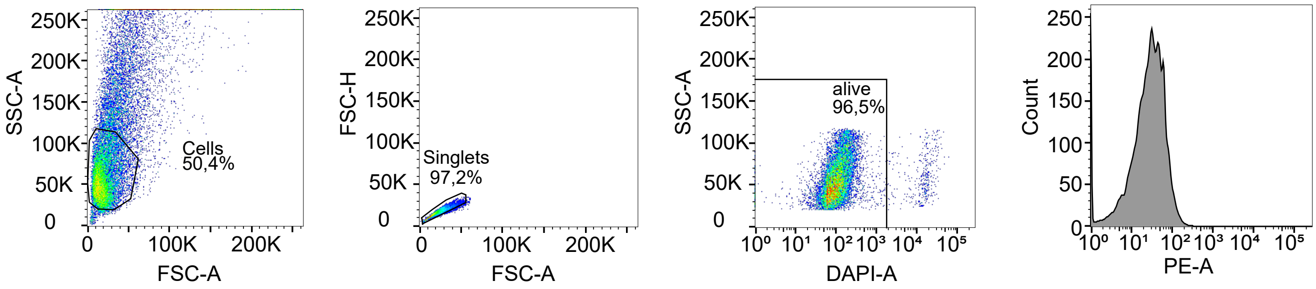


E

Mitochondrial membrane potential

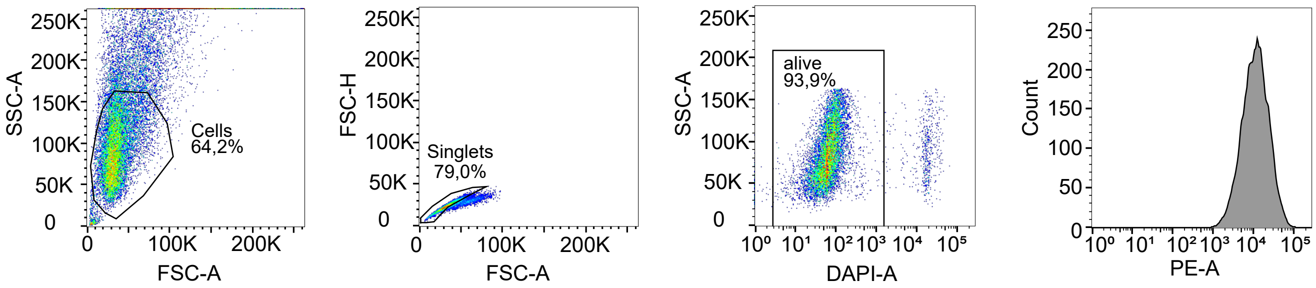

**Supplementary Figure 7. Representative flow cytometry gating strategies for cell barcoding, metastasis quantification, and metabolic assays. (A)** Gating strategy utilized to determine lentiviral barcode transduction efficiency and evaluate post-sorting purity. Sequential gating was performed as follows: initial cell populations were identified and debris excluded using forward scatter (FSC) versus side scatter (SSC) parameters, doublets were excluded via FSC-Area (FSC-A) versus FSC-Height (FSC-H) to isolate singlets, viable cells were selected using DAPI-Area (DAPI-A) versus SSC-A, and transduction efficiency was evaluated by quantifying the $GFP^+$ population via GFP-A versus SSC-A. **(B)** Gating strategy for the quantification of disseminated metastatic cells in cell suspensions from digested lung tissue. Following debris exclusion using FSC versus SSC, viable cells were identified via DAPI-A versus SSC-A, and the percentage of $GFP^+$ tumor cells was quantified using GFP-A versus SSC-A. **(C–E)** Gating strategies for functional mitochondrial and metabolic assays tracking **(C)** cytosolic ROS levels, **(D)** mitochondrial ROS (mtROS) levels, and **(E)** mitochondrial membrane potential (MMP). For these assays, sequential gating was performed to exclude debris (FSC vs. SSC) and doublets (isolating singlets via FSC-A vs. FSC-H), viable cells were selected via DAPI-A versus SSC-A, and target populations were quantified as Median Fluorescence Intensity (MFI) shifts in histogram plots using specific fluorophore channels (GFP, PE, or APC, depending on the cell line and assay configuration) relative to negative controls.
